# The *Pemphigus Vulgaris* antigen desmoglein-3 suppresses p53 function via the YAP-Hippo pathway

**DOI:** 10.1101/399980

**Authors:** Ambreen Rehman, Yang Cai, Christian Hünefeld, Hana Jedličková, Yunying Huang, M Teck Teh, Jutamas Uttagomol, Angray Kang, Gary Warnes, Usama Ahmad, Catherine Harwood, Daniele Bergamaschi, Eric Kenneth Parkinson, Martin Röcken, Ian Hart, Hong Wan

**Author notes:** Author contributions: H.W., A.R., Y.C., C.H., M.R., E.K.P. and D.B. designed research; H.W., I.H., A.R., J.U. and C.H. wrote the manuscript. A.R., H.W., Y.C., C.H., Y.H. and J.U. performed most of the experiments. M.T.T. performed qPCR; G.W. performed FACS; H.J., C.Ha., D.B., E.K.P., A.K. and U.A. contributed PV sera/cell line/reagents/analysis; H.W., A.R., Y.C., C.H., M.R., E.K.P., C.Ha., D.B., M.T.T., A.K. and G.W. analysed data. To whom correspondence should be addressed., **Dr. Hong Wan BDS, MSc, PhD**, Centre for Immunobiology and Regenerative Medicine, Institute of Dentistry, Barts and The London, School of Medicine and Dentistry, Turner Street, Whitechapel, London E1 2AD, ORCID iD: orcid.org/0000-0003-3794-5692.

## Abstract

Desmoglein-3 (Dsg3), the *Pemphigus Vulgaris (PV)* antigen (PVA), plays an essential role in keratinocyte cell-cell adhesion and regulates various signaling pathways implicated in the pathogenesis the PV blistering disease. We show here that expression of Dsg3 may directly influence p53, a key transcription factor governing the response to cellular stress. Dsg3 depletion caused increased p53 and apoptosis, an effect that was further enhanced by UV and mechanical strain and reversed by Dsg3 gain-of-function studies. Analysis in Dsg3-/- mouse skin confirmed increased p53/p21/caspase-3 compared to Dsg3+/- control *in vivo*. This Dsg3-p53 pathway involved YAP since Dsg3 forms a complex with YAP and regulates its expression and localization. Analysis of PV patient samples detected increased p53/YAP with diffuse cytoplasmic and/or nuclear staining in cells surrounding blisters. Treatment of keratinocytes with PV sera evoked pronounced p53/YAP expression. Collectively, our findings establish a novel role for Dsg3 as an anti-stress protein, via suppression of p53 function, suggesting that this pathway, involving YAP-Hippo control of skin homeostasis, is disrupted in PV.

## Introduction

Desmoglein-3 (Dsg3), a member of the cadherin superfamily, serves as an adhesion protein in desmosomes and is crucial in the maintenance of epithelia integrity. A growing body of evidence suggests that Dsg3 does not function solely in desmosome adhesion but also acts as a regulator for various signalling pathways that govern cell adhesion, proliferation, differentiation, morphogenesis, as well as cell migration (Brown and Wan, 2015; Chen et al., 2013; Mannan et al., 2011; Rotzer et al., 2016; Tsang et al., 2010; Tsang et al., 2012a; Tsang et al., 2012b). However, the non-junctional functions of Dsg3 remain poorly understood. Dsg3 is distributed on the cell surface beyond the desmosomes and, intriguingly, exhibits distinct tissue expression patterns between the skin and mucous membranes. While in the skin Dsg3 largely is restricted to keratinocyte basal and immediate suprabasal layers of the epidermis, in oral mucosa it exhibits uniform expression across all live stratified squamous epithelial layers (Amagai et al., 1996; Teh et al., 2011). Why the protein exhibits such a distinct tissue distribution pattern is unknown.

Dsg3 has been shown to be down regulated in the human autoimmune condition, *Pemphigus Vulgaris* (PV) and upregulated in squamous cell carcinoma. In PV, Dsg3 serves as a major antigen (PVA) and is targeted by autoantibodies, leading to steric hindrance of Dsg3 interactions followed by its depletion from desmosomes and the cell surface. As a result, disruption of cell-cell cohesion occurs that causes pemphigus acantholysis in the Dsg3 bearing tissues (Amagai et al., 1991; Calkins et al., 2006; Delva et al., 2008; Kitajima, 2014; Lanza et al., 2006). However, there is evidence that argues against this theory and suggests that PV is caused by antibodies targeting other surface receptors, rather than Dsg3, thereby triggering an intracellular signalling cascade, which leads to cell shrinkage, apoptosis and, ultimately, blistering (Amagai et al., 2006; Grando et al., 2009; Lanza et al., 2006). In consequence, the pathogenesis of PV remains not fully understood and constitutes an issue of debate (Amagai et al., 2006; Spindler et al., 2018). In cancer, Dsg3 is found to be upregulated, in particular in squamous cell carcinoma (Chen et al., 2007; Fang et al., 2014; Ferris et al., 2005; Fukuoka et al., 2007; Huang et al., 2010; Liu et al., 2007; Savci-Heijink et al., 2009); though its actual role in tumour progression and metastasis remains to be elucidated (Brown and Wan, 2015). *In vitro* experiments with gain of Dsg3 function support its pro-cancerous role, in line with the clinical findings, where overexpression of exogenous Dsg3 in cell lines elicits pronounced membrane protrusions and augments cell migratory capacity via activation of various signalling pathways, e.g. Src, Rac1/Cdc42, Ezrin and AP-1 transcription factor which are known to be ‘hijacked’ in cancers (Brown et al., 2014; Brown and Wan, 2015; Tsang et al., 2010; Tsang et al., 2012a; Tsang et al., 2012b). Conversely, Dsg3 depletion results in inhibition of tumour growth and metastasis (Chen et al., 2007).

Our preliminary observation, made in MDCK (Madin–Darby canine kidney) cells, showed that overexpression of Dsg3 resulted in suppression of dome formation, a phenomenon occurring spontaneously in prolonged MDCK cultures marking the initiation of epithelial cell differentiation (Lever, 1979). Many compounds known to induce cell differentiation are capable of provoking dome formation in MDCK (Kennedy and Lever, 1984; Leighton et al., 1970; Oberleithner et al., 1990). So we speculated that Dsg3 might exert a function in the suppression of epithelial cell differentiation. Indeed, staining for a senescence marker, beta-galactosidase (β-gal) showed a marked decrease of β-gal signals in Dsg3 overexpressing cells in contrast to control cells (data not shown). As p53 is a master regulator of cell differentiation, senescence, metabolism and apoptosis (Vousden and Lu, 2002), we decided to analyse p53 expression in these cell lines. Our preliminary data suggested that Dsg3 might play a role in governing the expression and function of p53 since the overexpression of Dsg3 resulted in a suppression of p53, and its downstream target p21^WAF1/CIP1^, as compared to empty vector control cells (data not shown). In the present report, we investigated the hypothesis that Dsg3 may negatively regulate p53 in keratinocytes subjected to UV irradiation and mechanical stress, and to explore its contribution to the pathogenesis of PV.

## Results

### Dsg3 depletion induces p53 expression and activity in keratinocytes

To investigate our hypothesis that Dsg3 may negatively regulate p53, in a physiologically relevant system, we performed an RNAi study in skin-derived NTERT keratinocytes that harbour wild type p53 (wtp53). Knockdown of Dsg3 caused no apparent changes in other junctional proteins in this cell line (Fig EV1A) so the effect of Dsg3 was not due to the mere disruption of keratinocyte intercellular adhesions (see below). However, immunofluorescence for p53 indicated a significant induction of nuclear p53 in the RNAi treated cells compared to controls transfected with scrambled siRNA (Fig 1A). Western blotting analysis detected only a moderate increase of p53 relative to control samples (Fig 1B). However p53 is a very dynamic protein (Purvis et al., 2012), so we asked whether this subtle change in steady-state p53 in Dsg3 depleted cells was masked by fast protein turnover. Hence, we treated the RNAi transfected cells with the proteasome inhibitor MG-132 (25 μM) for 3 hours before protein extraction; now a marked increase of p53, accompanied by p21^WAF1/CIP1^, was detected in Dsg3 knockdown cells (Fig 1C). To consolidate this finding, we monitored p53 protein turnover where siRNA transfected cells were treated with cycloheximide (30 μg/ml) before being harvested at various time points for up to 6 hours. As expected, only a delayed reduction of p53 was found in Dsg3-depleted cells such that by 6 hours, whereas p53 had almost disappeared in control cells, some residual p53 still was detectable in the knockdown cell samples. Calculation of p53 half-life indicated that p53 persisted at least 3-fold longer in Dsg3 knockdown cells than in control cells (>180min vs 65min), suggesting that Dsg3 depletion caused accumulation of p53 accompanied by stabilisation of MDM2 (Fig 1D), a key negative regulator of p53 (Kubbutat et al., 1997). We reasoned that this might be due to a negative auto-feedback loop between p53 and its targeted transcription regulation of the MDM2 gene (Piette et al., 1997). A time course analysis was performed and showed induction of p53 and p21^WAF1/CIP1^ after Dsg3 knockdown, accompanied by a loss (reduction) of MDM2 expression after 3 days of siRNA transfection (Fig EV1B). Analysis with the phospho-S166 antibody for MDM2 indicated enhanced phosphorylation of MDM2-S166 in knockdown cells suggesting the likelihood that this could increase cell survival by ubiquitining p53 and targeting it for degradation (Meek and Knippschild, 2003). Overall, these findings agree with our hypothesis that Dsg3 may restrain the p53 pathway, at least in part by modulating ubiquitin ligases such as MDM2.

**Figure 1.**
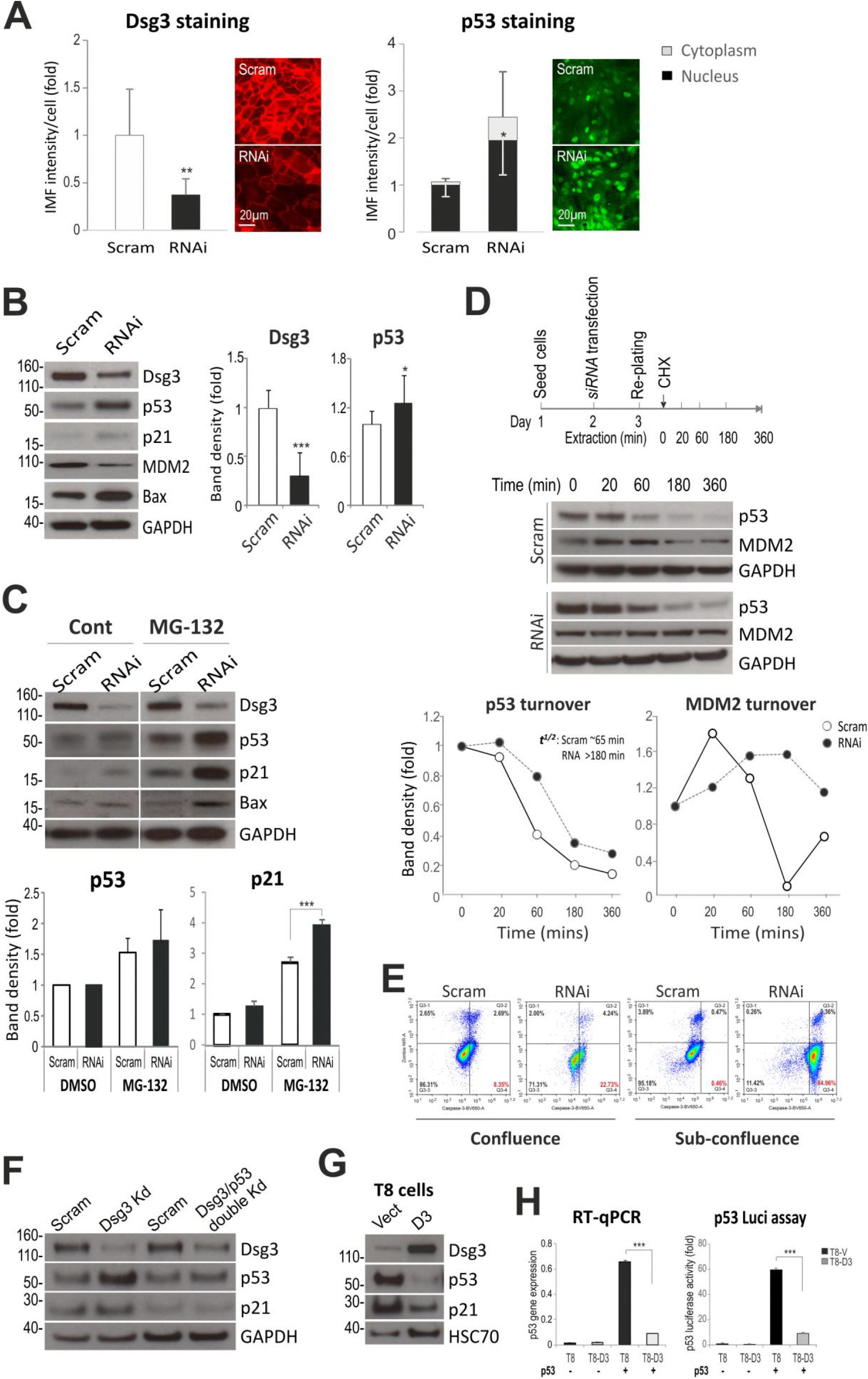
Dsg3 depletion in human keratinocytes enhances the p53 expression and activity. (**A**) Immunofluorescence analysis of NTERTs transiently transfected with Dsg3 specific or scrambled siRNA for 2d showed significantly increased nuclear p53 relative to control; n=7 fields/condition, pooled from 2 independent experiments. Scale bars, 20 μm. (**B**) Western blotting for p53 and its targets, p21^WAF1/CIP1^ and MDM2, in NTERTs with Dsg3 knockdown indicated a moderate but significant increase of p53 expression; n=4 biologically independent samples. (**C**) Western blots of siRNA-transfected cells with and without the treatment of MG-132 (25μM) for 3h. A marked increase of p53/p21^WAF1^/CIP^11^ was displayed in Dsg3 knockdown cells treated with MG-132 compared to control. GAPDH and HSC70 were used as loading controls. (**D**) Protein turnover analysis for p53 and MDM2 indicated reduced p53 turnover with concomitant MDM2 stabilisation in the Dsg3 depleted cells. The time line of the experiment was shown above the blots. The band density for each blot was normalised against the loading control in each sample and then against the one at 0 min time point in each condition. The calculated half-life for p53 was shown in the p53 graph. (**E**) Flow cytometric analysis of cell viability and apoptosis with live cells (Zombie NIR^-ve^/Caspase-3^-ve^), apoptotic (Zombie NIR^-ve^/Caspase-3^+ve^), late apoptotic (Zombie NIR^+ve^/Caspase-3^+ve^) and necrotic (Zombie NIR^+ve^/Caspase-3^-ve^) in NTERTs with and without Dsg3 knockdown, grown at 100% and ∼40% confluences; the represented data of 3 independent attempts. (**F**) Western blots with the indicated antibodies in lysates of cells with single (Dsg3) and double (Dsg3/p53) knockdown for 2d. (**G**) Western blots of stable cutaneous keratinocyte T8 Vect control and Dsg3 overexpression (D3) (p53 null) lines showed suppression of p53/p21^WAF1/CIP1^ in D3 cells compared to Vect cells. (**H**) Left: RT-qPCR analysis of p53 expression (mean±s.e.m.) in T8 cell lines with or without p53 transfection (n=3 independent assays of duplicate in each test). Right: p53 luciferase assay (mean±s.d.) of T8 cell lines with and without p53 transfection (n=3, a representative of two independent experiments). Comparison was via unpaired two-sided student *t*-test. *p<0.05, ***p<0.01.

To further define the apoptotic phenotype, we performed the FACS based Zombie NIR-caspase-3 assay (Lee et al., 2018) in siRNA pre-treated cells grown to confluent and subconfluent conditions and detected a marked increase of classic apoptotic cells (Zombie NIR^-ve^/Caspase-3^+ve^) in Dsg3 knockdown cells as compared to the respective controls (Fig 1E). Moreover, the induction of p53/p21^WAF1/CIP1^ caused by Dsg3 depletion was attenuated with double knockdown in cells for Dsg3 and p53 (Fig 1F). Finally, we evaluated our finding in another cutaneous cell line, T8 (that harbours a frameshift mutation at amino acid 91 of TP53 resulting in a truncated protein and thus it is essentially null in p53) with transduction of hDsg3.myc. After transient transfection of wtp53 plasmid for two days, we observed marked suppression of p53, compared to empty vector cells, at both the protein and gene levels; this was further confirmed by the p53 luciferase assay (Fig 1G,H).

To explore whether this pathway has any influence on the p53 mediated regulation of cell differentiation, we performed qPCR analysis for various genes involved in early and late differentiation programmes in keratinocyte lines with modulation of Dsg3 expression levels, i.e. Dsg3 knockdown or ectopic overexpression. In this approach, we found a general inverse relationship between Dsg3 expression and key differentiation markers of keratinocytes (Supplemental Figure 2). Dsg3 silencing caused enhanced expression of terminally differentiating structural genes indicating a premature cellular differentiation, while the inverse result was seen in cells with Dsg3 overexpression indicating cell dedifferentiation. These results suggest that the Dsg3-p53 pathway also has some influence, at least in part, on the cell differentiation programme regulated by p53. Alternatively, the results seen in Dsg3 loss-of-function cells may be related to the induction of keratinocyte senescence that culminates in increased differentiation. Indeed preliminary data from the Dsg3-/- mice showed increased p16^INK4A^ staining in the skin, supporting this hypothesis.

**Figure 2.**
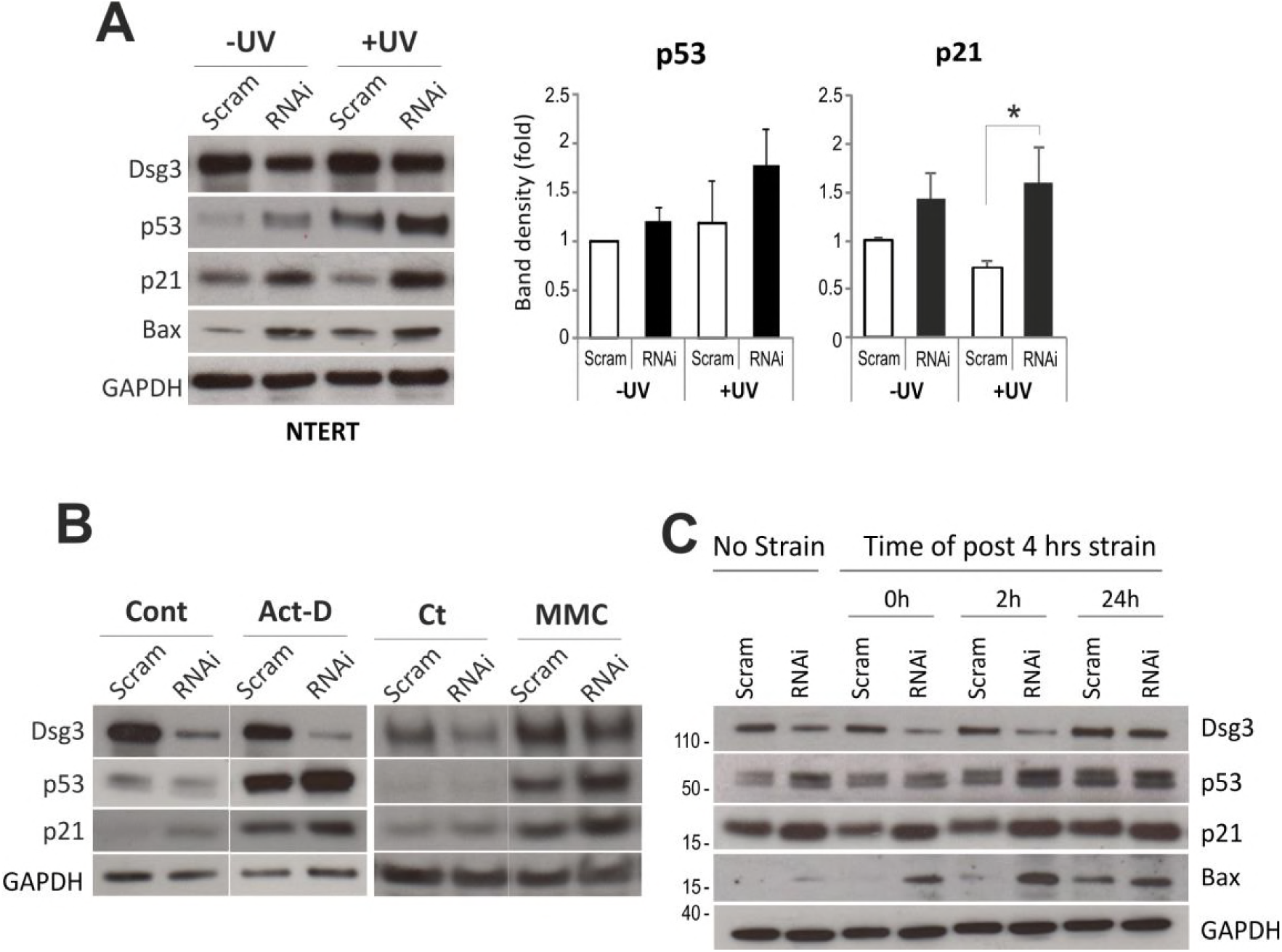
Dsg3 depletion causes further induction of the p53 expression and activity in response to stress signals. (**A**) Western blotting analysis of siRNA pre-treated NTERT cells with and without UVB irradiation for the indicated proteins (n=3 biologically independent samples, *p<0.05). (**B**) Western blotting analyses for the p53 and p21^WAF1/CIP1^ expression in cells treated with or without actinomycin D (Act D, 5nM) and mitomycin (MMC, 5ug/ml) for 24h, respectively. Enhanced expression of p53 and p21^WAF1/CIP1^ was shown in cells treated with drugs in control cells compared to cells without the drug treatment, but with further induction shown in Dsg3 depleted cells exposed to drugs. (**C**) Mechanical loading of NTERTs with cyclic strain caused further induction of p53, p21^WAF1/CIP1^ and Bax in cells with Dsg3 knockdown relative to the respective controls. The siRNA pre-treated cells were seeded at confluent density in BioFlex plates after transfection, and then subjected to cyclic strain (TX-5000, 1Hz, 20% amplitude) for 4 hours the following day. Lysates were extracted either immediately after strain or 2h and 24h later, respectively, after transference of the plates to a stationary state in an incubator, along with static control cells.

To determine whether the pathway of Dsg3-p53 is Dsg3-specific we performed similar knockdown experiments for desmoplakin, a marker of desmosomes and E-cadherin, a classical cadherin in adherens junctions, in NTERTs and found that neither desmoplakin nor E-cadherin depletion resulted in comparable effects (Fig EV3A,B). Only a small reduction of p53, and its downstream targets p21^WAF1/CIP1^ and Bax, was detected in cells with E-cadherin knockdown and this was further enhanced by UV irradiation. Similarly, only a slight reduction of p53 was shown in cells with desmoplakin knockdown. It had no effect on p21^WAF1/CIP1^ and marginally increased Bax. Collectively, these data suggest that the negative regulation of p53 by Dsg3 likely is independent of the desmosomes and adherens junctions, suggesting that such a function might be attributed to the extra-junctional pool of Dsg3 (Spindler et al., 2018).

### Dsg3 antagonises the p53 expression and function in response of keratinocytes to stress signals

The Dsg3 expressing tissues, such as skin and oral mucosa, are exposed daily to various chemical insults and physical stresses that, in theory, are capable of triggering p53 induction because of its central role in response to cellular stress signals (Vousden and Lu, 2002). To delineate the contribution of Dsg3 in such cellular responses, we decided to challenge the siRNA treated NTERTs with various stresses, such as UV exposure and mechanical stretching, before analysis of p53 expression. We asked whether Dsg3 has the ability to suppress or counterbalance the p53 response to these stress signals. First, we analysed cells subjected to UVB irradiation (10∼30mJ/cm^2^) and observed a trend of elevated p53 accompanied with a significant increase of p21^WAF1/CIP1^ after one day. This effect was significantly enhanced in Dsg3 depleted cells as compared to controls, indicating that Dsg3 has the ability to antagonise UV induced p53 expression (Fig 2A). Consistently, we observed that overexpression of Dsg3 resulted in suppression of p53 and p21^WAF1/CIP1^ after UV exposure (Fig EV4A). Moreover, we found that cells overexpressing Dsg3 were highly resistant to cell death induced by UV; in contrast, control cells were highly sensitive to UV-induced cell death. Similar findings were made with another two cell lines A2780 and HCT116 (wtp53) where transduction of Dsg3 protected cells from UV-induced death, as compared to Vect control cells (Fig EV4B). In line with this, cells treated with genotoxic drugs, such as actinomycin D or mitomycin C, also showed a similar effect with a strong induction of p53 and p21^WAF1/CIP1^ in Dsg3 depleted cells as compared to cells without Dsg3 knockdown or with knockdown but without drug treatment (Fig 2B). Together, these data suggest that Dsg3-reduced modulation of stress responses is a general phenomenon and that overexpression of Dsg3 protects cells against various environmental insults by dampening the p53 response.

Next, we challenged either siRNA treated cells or control cells, with mechanical loading using equiaxial cyclic strain (FX-5000, 1Hz, 20%) for 4 hours. Lysates were extracted immediately after strain (0h) or 2 and 24 hours later, respectively. As shown in Figure 2C, increased p53 and p21^WAF1/CIP1^/Bax were detected in Dsg3 knockdown cells, in particular at 2h and 24h time points in post-strained cells, compared to the respective controls. This result indicated that the loss of Dsg3 function also affected p53 stabilisation and activity in response to mechanical stress, reinforcing the ability of Dsg3 to counterbalance the p53 response to mechanical stress.

### Increased p53 expression and apoptosis is detected in Dsg3-/- mouse skin in vivo

Having confirmed the pathway of Dsg3 negative regulation of p53 in keratinocytes, we then wished to determine whether alteration of this pathway is detectable in Dsg3 knockout mice. Mice with a targeted ablation of Dsg3 develop a phenotype of runting and wave-pattern hair loss accompanied with oral and skin lesions after weaning (Koch et al., 1997; Koch et al., 1998). Hence, we evaluated the expression of p53, p21^WAF1/CIP1^ and cleaved caspase-3 in the back skin tissue samples at 8∼12 weeks and compared the finding with that of healthy heterozygous littermates (Dsg3+/- control). Increased expression of p53, p21^WAF1/CIP1^ and cleaved caspase-3 proteins was detected in Dsg3-/- hair follicles but not in Dsg3+/- controls (Fig 3). This result confirmed that Dsg3 expression prevents p53 activation and apoptosis in mouse skin and suggests that the phenotype of Dsg3 null mice may result, at least in part, from accelerated and/or enhanced keratinocyte death.

**Figure 3.**
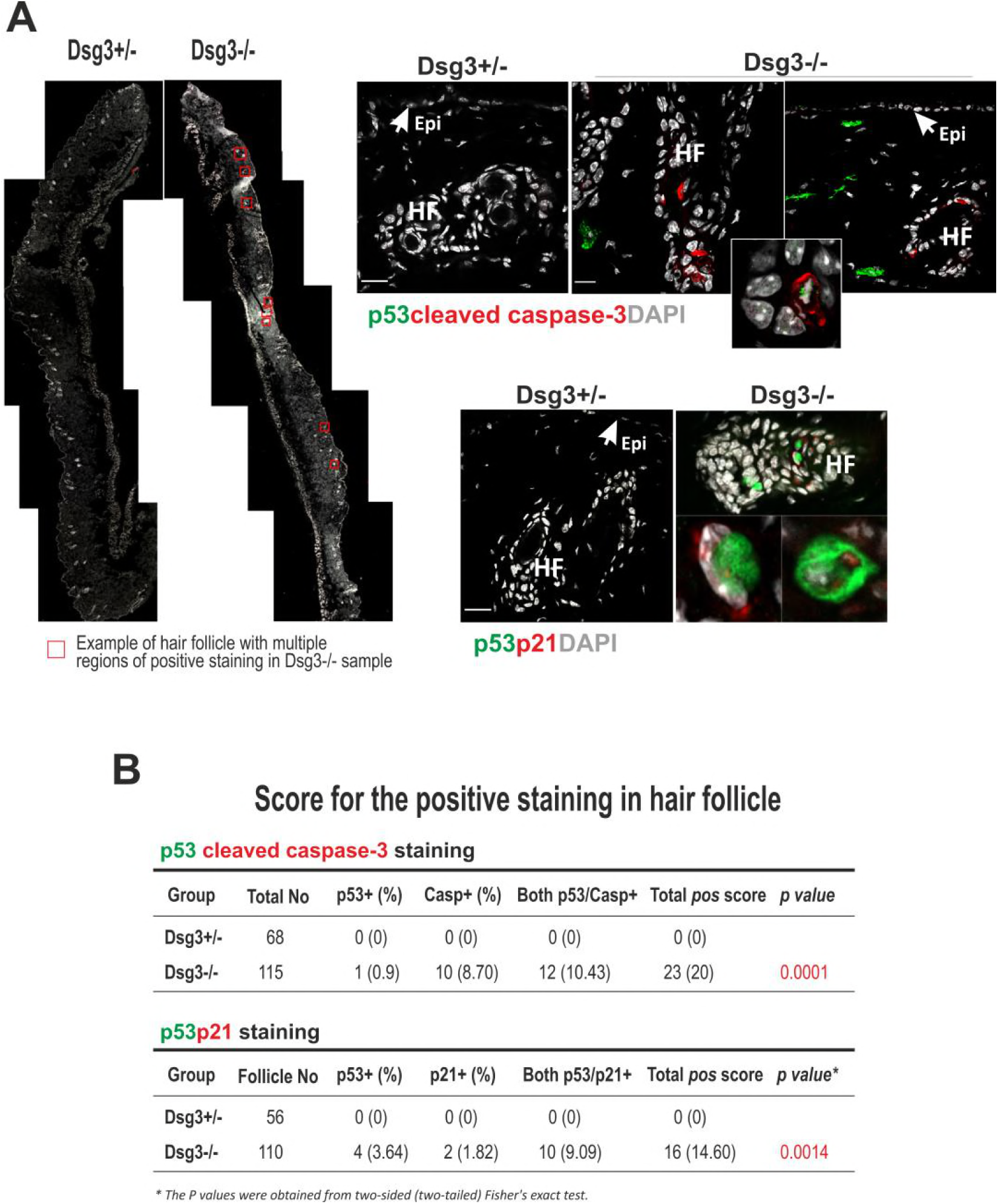
Increased expression of p53, p21^WAF1/CIP1^ and cleaved caspase-3 is observed in the back skin of Dsg3 knockout mice. (**A**) Immunofluorescent staining in the back skin of Dsg3-/- and Dsg3+/- (heterozygous littermate) mice showed elevated staining signals for the indicated proteins in the hair follicles of Dsg3-/- mice as compared to heterozygous littermate, though no positive staining was yet observed in the epidermis (n=2 mice per group, aged 8∼12 weeks). Some fibroblasts in the dermis were also shown positive staining of p53. Epi: epidermis, HF: hair follicle. The insert in the top right panels highlights cells with double positive staining for p53 and active caspase-3 in Dsg3 null skin. Scale bar, 20 μm. (**B**) Tables summarise the scores of positive hair follicle staining for p53/active caspase-3 and p53/p21^WAF1/CIP1^, respectively. Each hair follicle containing one or more positively stained keratinocytes was scored positive.

### Dsg3 forms a complex with YAP and regulates YAP function

It has been shown that the p53 response to DNA damage is attenuated by high cell density in which the Hippo pathway is activated (Aqeilan, 2013; Bar et al., 2004; Reuven et al., 2013). However, this mechanism has not been thoroughly investigated in human keratinocytes. Given that, 1) the biological functions of p53 and the Hippo pathway overlap at multiple levels and their upstream cues remain largely unknown (Furth et al., 2018); 2) the Hippo pathway is required in E-cadherin mediated contact inhibition of proliferation (CIP) in dense cell populations (Kim et al., 2011) and as we and others have shown that Dsg3 engages in cross talk with E-cadherin and regulates its adhesive function (Rotzer et al., 2015; Tsang et al., 2010; Tsang et al., 2012b), we decided to analyse the Hippo pathway and its downstream effector, YAP (Furth et al., 2018). First, we analysed, by Western blotting, the expression of YAP, as well as pYAP-S127 (an indicator of the Hippo pathway (Piccolo et al., 2014)), alongside p53 in confluent NTERT cells with and without Dsg3 knockdown and subjected the cells to equiaxial cyclic strain (the experiment as described above). We showed that in control cells, both Dsg3 and p53 exhibited a response to mechanical loading with Dsg3 induction and p53 suppression. The expression of YAP also showed alterations with moderate decline over time that reached to the lowest level at 24 hours in post-strained cells (Fig 4A). The expression of pYAP-S127 showed a similar trend, indicating that the Hippo pathway was perturbed temporally by mechanical stretching. Significantly, we observed evident reduction of YAP as well as pYAP in the Dsg3 knockdown cells, compared to controls, in both stationary and strained cells (no strain and 0h of post-strain Figure 4A). The residual Dsg3 in the knockdown cells also exhibited a response to mechanical strain by showing a steady increase up to 24 hours post-application of the stimulus. Collectively, these results indicate that mechanical loading evokes coordinated increases in Dsg3 levels with decreases in p53 and YAP levels.

**Figure 4.**
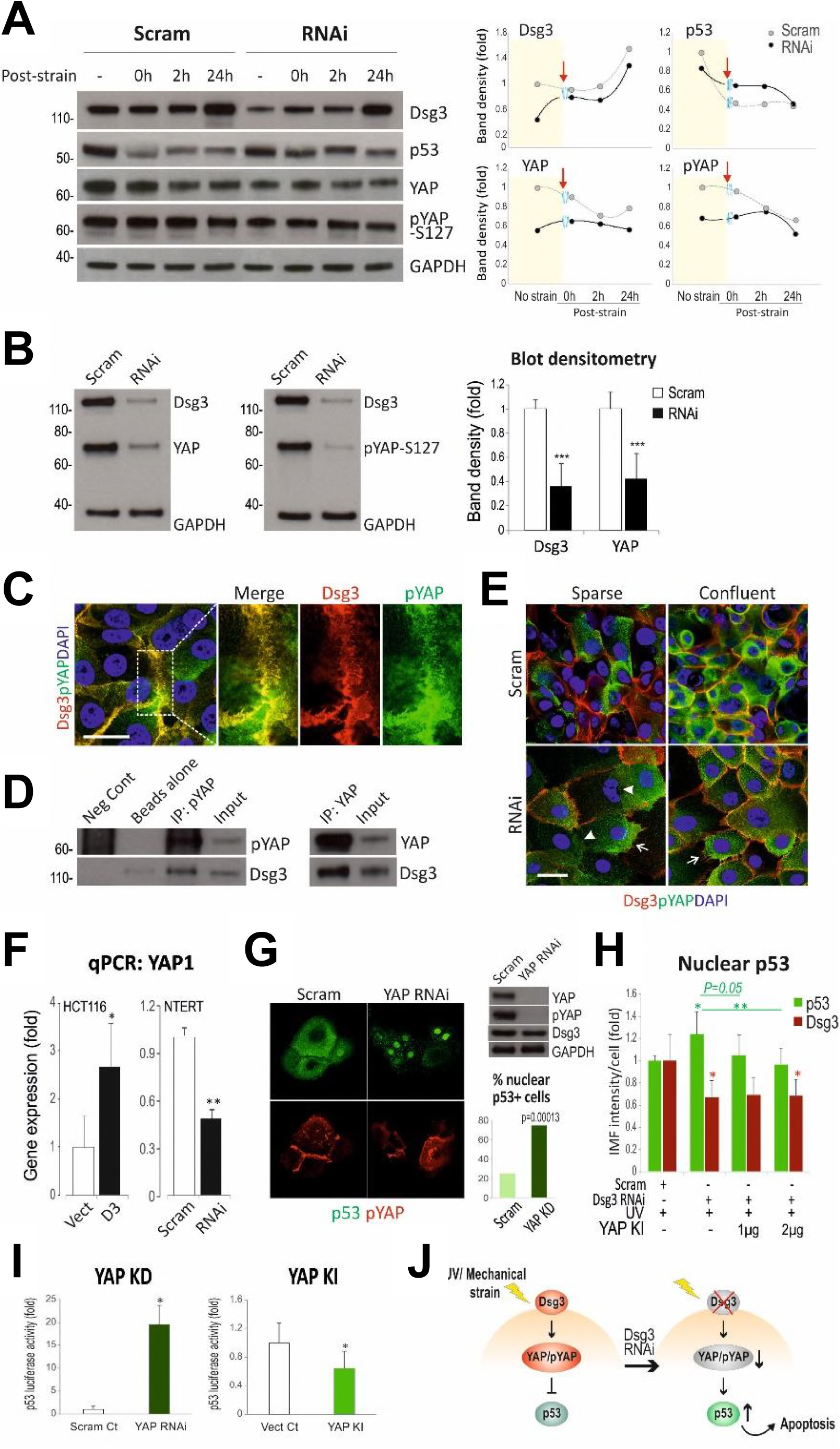
Dsg3 forms a complex with YAP and regulates its expression. (**A**) Western blotting for the indicated proteins in cyclic strained and non-strained cells. The densitometry analysis for each protein with a trend change is displayed on the right; light yellow areas indicated data of non-strained cells and the red arrows indicated the 0h points of post-strained cells. (**B**) Effect of Dsg3 knockdown on YAP and pYAP in steady state that showed significant reduction compared to controls (n=3 biologically independent samples per condition). (**C**) Super resolution microscopy of NTERT cells seeded at confluent density and dual stained for Dsg3 and pYAP revealed colocalisation of two proteins. The enlarged insert for each channel is shown on right. (**D**) Co-immunoprecipitation with antibodies for pYAP and YAP demonstrated that Dsg3 physically interacted with pYAP and YAP. (**E**) Confocal images of siRNA pre-treated NTERT cells seeded at low and high densities, and immune-labelled for Dsg3 and pYAP (arrowheads indicate the loss of both Dsg3 and pYAP at the cell periphery, whereas arrows indicate a trend of the pYAP endocytosis from the membrane where limited Dsg3 was present in knockdown cells). Scale bars in C&E, 10 μm. (**F**) qPCR analysis of the YAP gene expression in cells with Dsg3 overexpression or knockdown (n=3 biologically independent samples per condition). (**G**) Immunofluorescence of NTERTs without or with YAP knockdown, and labelled for p53 and pYAP. Cells seeded at low density in low calcium medium overnight before being transferred to high calcium keratinocyte growth medium (KGM) for 6h. Increased nuclear p53 was detected in the population of cells with YAP knockdown, and the score for the percentage of positive nuclear p53 was shown in the bar chart (n=6 fields per condition). Western blots of YAP and pYAP are displayed on the top. (**H**) p53 luciferase assay in cells with YAP knockdown or knock-in (n=3 biological independent samples). (**I**) Image quantitation data for the positive p53 nuclear expression in siRNA pre-treated NTERTs transfected with or without YAP plasmid at different concentrations. Cells were transfected with siRNA for 1 day and then with YAP plasmid for 5 hours before harvesting and seeded on coverslips. The next day, they were subjected to UV irradiation and fixed 6h later (n=6 fields per condition, data were mean±s.d.). All statistical comparisons were via unpaired two-sided Student’s *t*-test. *p<0.05, **p<0.01***p<0.001. (**J**) Schematic model of Dsg3 regulation of YAP/pYAP and suppression of p53 in response to environmental stress signals.

To confirm the finding that Dsg3 depletion caused a reduction of YAP, we repeated Western blotting of steady-state cells with and without Dsg3 knockdown. Consistently, a marked reduction of YAP/pYAP was detected in knockdown cells compared to controls (Fig 4B). Significantly, we observed that membrane distribution of pYAP in NTERTs, and to a lesser degree for YAP, colocalised with Dsg3 (Fig 4C). Co-immunoprecipitation assays performed using lysates extracted from freshly confluent cells confirmed that both pYAP/YAP were interacting physically with Dsg3 (Fig 4D), suggesting the likelihood of their functional association with each other. Loss of Dsg3 seemed to abolish membrane localisation of pYAP, in particular in cells seeded at low cell densities (arrowheads Figure 4E). Concomitantly, the endocytic processing of pYAP was noticeable in areas adjacent to the plasma membrane where limited Dsg3 was present (arrows Figure 4E). Significantly, we found that the regulation of YAP by Dsg3 also occurred at the mRNA levels since overexpression of exogenous Dsg3 caused marked increase of YAP1 whereas the opposite pertained in Dsg3 knockdown cells (Fig 4F). These results suggested that the regulation of YAP by Dsg3 might be related to the presence of wild type p53. Next, we monitored YAP/pYAP expression, with respect to Dsg3 in various maturation ages of confluent cell cultures grown in calcium medium for up to 8 days, by fluorescent microscopy and found that their expression and the membrane colocalisation between Dsg3 and pYAP altered as a function of time (Fig EV6A,B). In freshly confluent populations strong signals were detected (6 h∼3d). After 3 days, levels of both Dsg3/pYAP and their membrane distribution were falling and by day 6, the membrane distribution of pYAP became internalised; results which are consistent with Hippo turning off when the cell junctions were well established (see below). In line with the above hypothesis, Western blotting analysis showed similar expression profile for pYAP albeit the total YAP was relatively consistent over this period (Fig EV6C). Collectively, these findings establish that Dsg3 has a role in the positive regulation of YAP at both the transcriptional and translational levels and suggest a substantial role for Dsg3 in the regulation of the Hippo pathway in keratinocytes.

The specificity of Dsg3 in regulating the YAP-Hippo pathway also was addressed by analysing YAP/pYAP expression in NTERTs with either desmoplakin or E-cadherin knockdown, in conjunction with UV irradiation (Fig EV3C). While desmoplakin knockdown resulted in a small induction of YAP/pYAP, which was less in cells treated with UV, little or no changes were detected in cells with E-cadherin knockdown. Thus, the regulation of YAP/pYAP by Dsg3 appeared independent of desmosomes and adherens junctions, similar to our findings for the Dsg3-p53 pathway.

### YAP and p53 antagonise one another, downstream of Dsg3, in keratinocytes

Having established a role for Dsg3 in regulating the YAP-Hippo and p53 pathways in keratinocytes, respectively, we hypothesised that the negative regulation of p53 by Dsg3 may be through a mechanism involving Hippo signalling by sequestering pYAP to the cell surface; this, in turn, could inhibit p53 nuclear transcription function (working model: Dsg3->Hippo-|p53). Thus, we anticipated that depletion of Dsg3 could result in Hippo failure leading to an induction of p53. To test this possibility, we investigated the impact of YAP knockdown, as well as knock-in, on p53 expression (downstream of Dsg3), with a focus on p53 nuclear localisation. First, we performed YAP knockdown with siRNAi in NTERTs. Since YAP is inactivated at high cell density when Hippo is activated, we examined cells seeded on coverslips at low density. After 6 hours incubation in keratinocyte growth medium, coverslips were fixed and processed for immunostaining for p53/pYAP and YAP/Dsg3, respectively. We found that knockdown of YAP resulted in cell morphological changes with many cells appearing larger in size (Fig EV7) that phenocopied the changes observed with Dsg3 knockdown (Brown et al., 2014). Moreover, YAP knockdown impacted on Dsg3 junctional assembly though no evident change was shown in Dsg3 levels. Nuclear p53 staining was observed in ∼25% cells in control population, in particular at early stages of colony development (Fig 4G). In contrast, YAP knockdown resulted in significant increased nuclear p53 with ∼75% cells showing nuclear p53. Apparently, knocking down YAP, with concomitant Hippo disruption, evoked p53 nuclear accumulation that likely triggers cell cycle arrest or death. Indeed, the p53 luciferase assay demonstrated that YAP knockdown caused enhanced p53 promotor activity and an inverse effect was detected in cells with ectopic YAP overexpression (knock-in) indicating an antagonistic relationship between the YAP and p53 pathways in keratinocytes (Fig 4H). Hence, YAP depletion causes p53 induction, at the protein and transcriptional levels as well as its nuclear relocation in keratinocytes.

Next, we performed a rescue experiment with YAP knock-in in NTERT cells with Dsg3 depletion in conjunction with UV irradiation. To this end, the siRNA pre-treated cells were subjected to transfection with the YAP plasmid at different concentrations the next day prior to harvesting and re-plated on coverslips at low density. After overnight culture, all samples were subject to UV irradiation (10∼30 mJ/cm^2^) and then incubated in keratinocyte growth medium for 6 hours before immunostaining for p53 and Dsg3. The result showed that the increased expression of nuclear p53 in Dsg3 knockdown cells was significantly attenuated in cells with YAP knock-in, in particular in cells treated with higher concentration of the YAP plasmid, indicating that exogenous expression of YAP was capable of rescuing the UV-induced augmented p53 and activity (Fig 4I). Notably, the colonies appeared larger in the population of cells with ectopic expression of the YAP gene. These findings suggest that overexpression of YAP is able to block the UV-induced p53 expression and activity to some extent thereby improving cell survival in keratinocytes.

### Enhanced p53 and YAP expression is detected in PV in vivo as well as in keratinocyte cultures treated with PV sera in vitro

Compelling evidence suggests a fundamental role for Dsg3 in pemphigus pathogenesis (Amagai et al., 2006; Kitajima, 2014; Spindler et al., 2018). However, the exact mechanisms underlying pemphigus acantholysis remain poorly understood and still are under debate in the field (Amagai et al., 2006; Spindler et al., 2018). *In vitro* studies demonstrate clearly that treating cells with autoantibodies from PV patients causes depletion of Dsg3 from desmosomes and the cell surface, leading to disruption of cell-cell junctions and collapse of the keratinocytes (Calkins et al., 2006; Delva et al., 2008). To explore whether the pathway suggested by our *in vitro* studies is operative in PV we performed immunohistochemistry for p53, as well as YAP, in oral mucous tissue biopsies from 25 PV patients. We found enhanced positive p53 staining in both cytoplasm and nucleus in at least 12 of 25 PV cases, especially in cells surrounding or in the clusters within blisters. In contrast, the normal samples showed only a few cells, located in the basal and suprabasal layers, with positive nuclear p53 staining (Fig 5A). Immunostaining for the active caspase-3 in PV samples with positive p53 also showed positive staining of caspase-3, confirming cell apoptosis in PV (Fig EV8). The p53 staining in PV was quite different from that observed in oral cancer, which showed strong nuclear signals, likely reflecting mutant p53 staining. YAP staining in PV exhibited more variations, with enhanced cytoplasmic staining in some cases (5 of 15 YAP positive staining) and pronounced nuclear YAP in others (10 of 15 positive staining) compared to the almost negative control (Fig 5B). By contrast, oral cancer showed augmented positive staining in both the nucleus and cytoplasm. Within the 25 PV cases, 8 were shown positive and 6 were negative for staining for both proteins, 4 were positive only for p53 and 7 were positive only for YAP, regardless of their locations. Furthermore, positive staining for both proteins was also observed in non-lesional areas in PV (Case3 in Figure 5A,B). These results indicate clearly alterations in the pathway of Dsg3-Hippo-p53 in PV that likely contributed to the pathogenesis of blistering in this disease.

**Figure 5.**
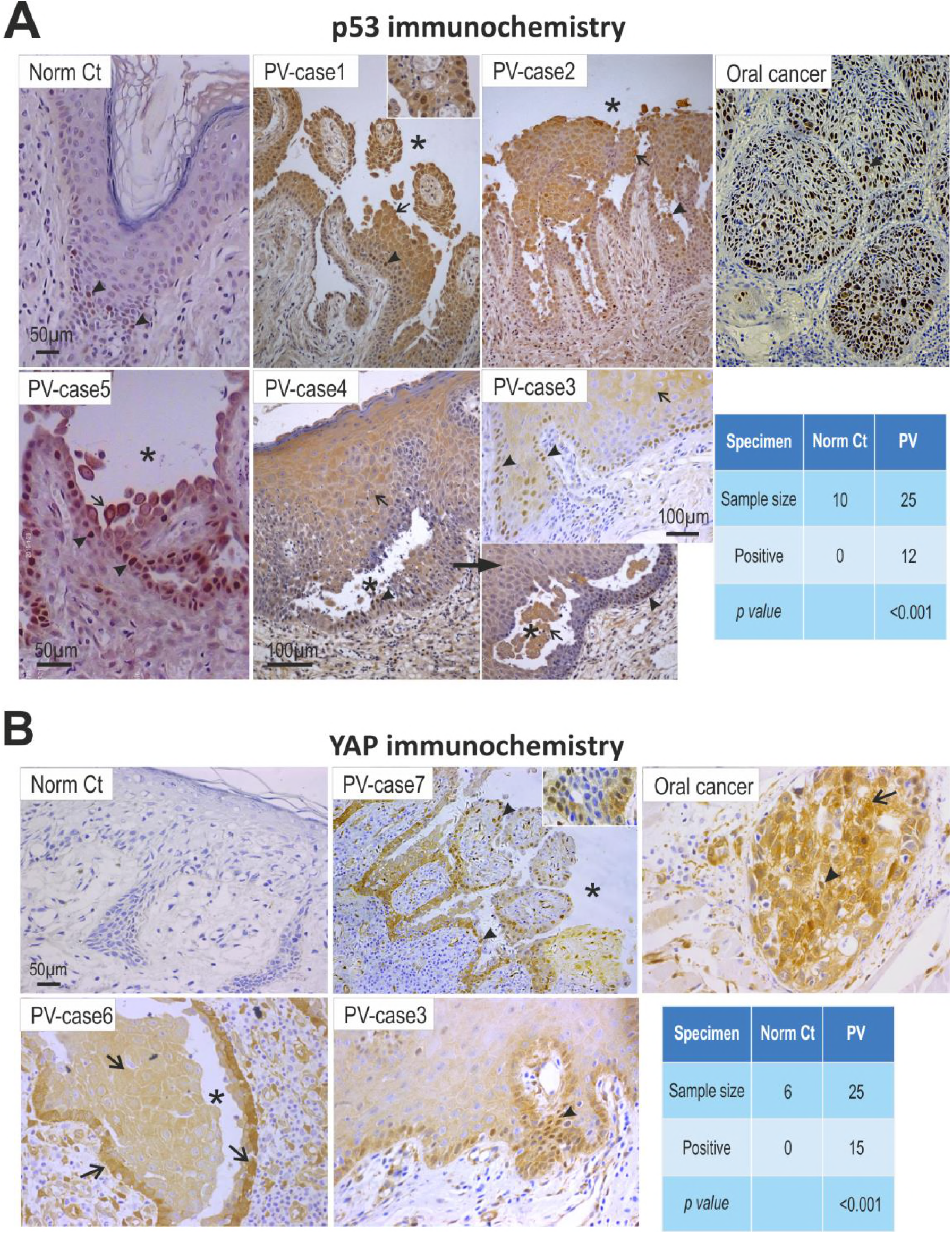
Enhanced p53 and YAP expression is shown in clinical PV patient samples and also in keratinocyte cultures treated with PV sera. p53 (**A**) and YAP (**B**) immunohistochemistry in oral mucous tissues from PV patients, and significant enhanced p53 staining was detected in 48% patients (arrowheads indicated positive nuclear staining whereas arrows indicated predominant cytoplasmic staining) and increased YAP either in cytoplasm (arrows) or nucleus (arrowheads), respectively, in 60% patients compared to normal controls. Oral mucous cancer was used as the positive control here. Asterisks indicate the areas of the blisters. Scale bars, 50 μm.

To validate the clinical findings, we next performed an *in vitro* study with PV sera collected from a different cohort of 17 patients as well as with a well-characterised pathogenic monoclonal antibody, AK23, that targets the adhesion site at the N-terminus of Dsg3 (Tsunoda et al., 2003). NTERT cells seeded at confluent densities were treated with PV sera or AK23 at various concentrations (40% PV sera, 1∼100μg/ml AK23) for different periods before immunostaining for p53, YAP and Dsg3. The result showed that, in control cells treated with normal sera, nuclear p53 was predominant, with limited cytoplasmic staining. By contrast, cells treated with PV sera (for 24 hours) exhibited remarkable augmentation in cytoplasmic p53 although there were variations between different patient samples (Fig 6A-C). Whist in some PV serum treated-cells membranous and cytoplasmic staining of p53 was evident others exhibited drastic reduction of Dsg3 and its disruption at the junctions (arrowhead in Figure 6A right panel). Intriguingly, the membrane distribution of p53 showed colocalisation with Dsg3 when there was severe membrane disruption (Fig 6A arrows in the inserts). The Dsg3 staining, however, exhibited broad variations and not all samples treated with PV sera (24 hours) showed Dsg3 depletion from the cell surface (Fig 6B). Indeed, some PV serum treated samples even showed a marked increase accompanied with pronounced Dsg3 disruption at the junctions and its aggregates in the cytoplasm (eg. PV serum-12, Figure 6B).

**Figure 6.**
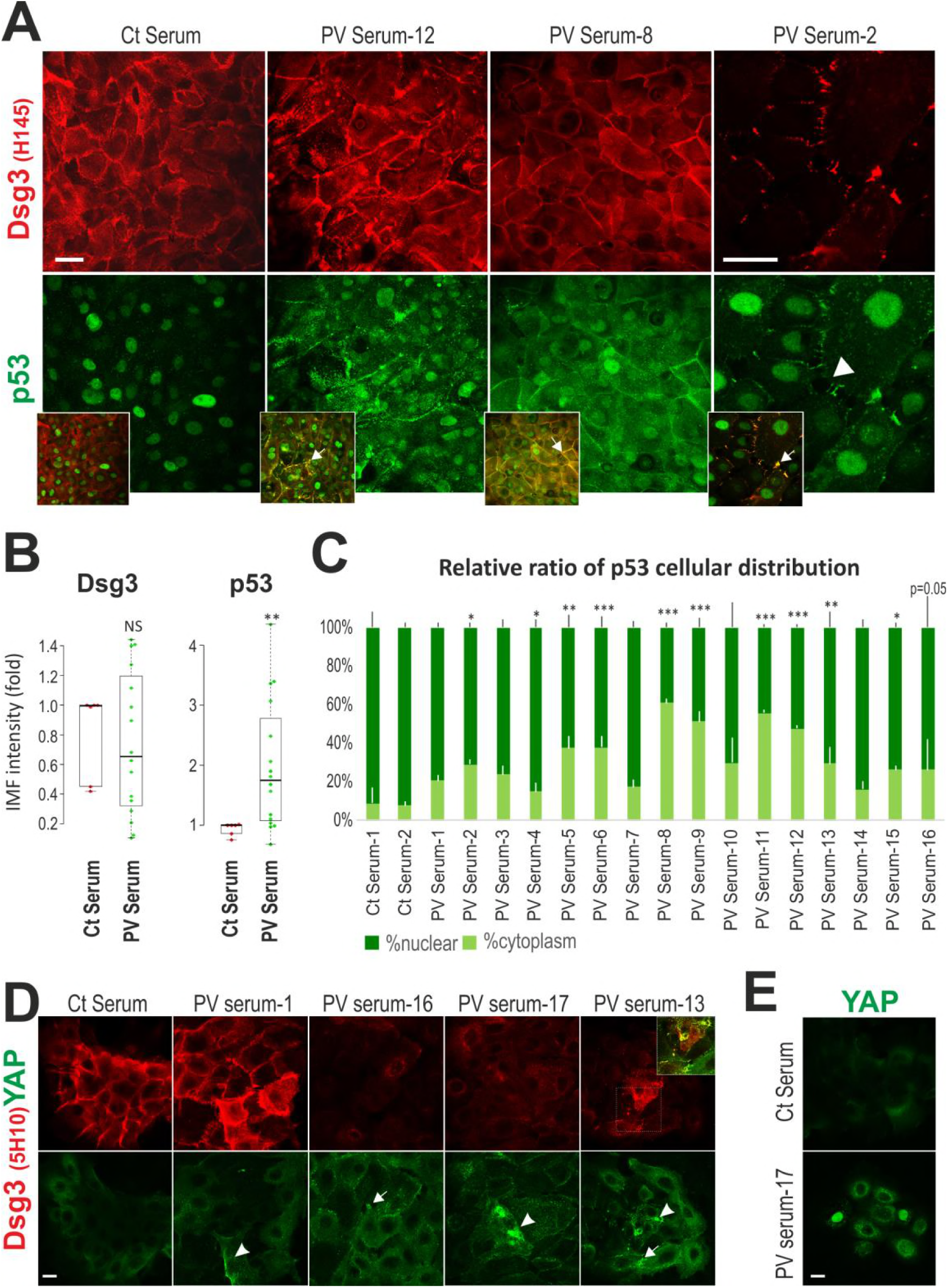
Altered p53 and YAP expression and distribution were also detected in keratinocyte cultures treated with PV sara. (**A**) Confocal microscopy of NTERT cells dual labelled for Dsg3 and p53. Cells were seeded at confluent densities in KGM for overnight before being treated with PV sera (at 40% concentration in KGM) from a different cohort of PV patients (n=17), for 24 hours before immunostaining for the indicated proteins. Disruption or depletion of Dsg3 at the plasma membrane accompanied with marked increases in p53 were observed in PV serum treated cells compared to controls exposed to sera of health individual that displayed, predominantly, nuclear p53 signals. Of note, p53 also showed distributed at the membrane where it colocalised with the fragmented Dsg3 (arrows). (**B**) Scatter and whisker plots of the Dsg3 and p53 cellular expression (n=16 for PV serum samples, n= 6 for control samples). Significantly increased nuclear p53 was also detected in PV serum treated cells (data not shown). (**C**) Relative ratio of p53 nuclear versus cytoplasmic cellular distribution in controls and 16 PV sera treated NTERTs. (**D**) The representative confocal images of NTERTs dual labelled for Dsg3 (5H10) and YAP. Cells seeded at low density were treated with PV sera (at 40% concentration in KGM) for 24 hours. A correlation between the depletion of Dsg3 and increase of YAP in the cytoplasm (arrowheads) as well as at the plasma membrane (arrows) was shown. Colocalisation of Dsg3 and YAP was highlighted in the insert. (**E**) Nuclear YAP staining was detected in PV serum treated cells seeded at clonal density in contrast to control serum treated cells that presented weak diffuse cytoplasmic staining. Chi-Square statistic was used to obtain the p values in A & B; Student *t*-test and the Wilcoxon-Mann-Whitney Rank Test were used for statistical significant analysis in D and gave similar results. *p<0.05, **p<0.01, ***p<0.001. NS: not significant.

Accompanying these changes in p53, the YAP staining also showed membrane disruption and concomitant cytoplasmic accumulation in the PV sera treated cells, with a clear correlation between the degree of Dsg3 loss (labelled with 5H10) and positive YAP staining (p<0.05, Figure 6D), suggesting the YAP accumulation owing to the degeneration of the Hippo pathway. (It is worth noting that cells displaying the negative staining with the monoclonal antibody 5H10 were labelled positive with H145, a rabbit antibody that binds to the cytoplasmic tail of Dsg3.) In contrast, the control serum treated cells showed only weak YAP staining in dense cells. These results seemed to recapitulate the observation in some PV samples (see Figure 5B). In addition, we also observed that the control serum treated cells, seeded at low density, presented a tendency of cell stratification in the colonies. On the other hand, the PV sera treated cells displayed a trait of flattened morphology with remarkable disruption of YAP at the plasma membrane, indicating perturbation of junction formation and cell polarisation. Furthermore, in smaller colonies containing a few cells in which the junctions were less mature, the nuclear YAP was detectable (Fig 6E). This, again, recapitulated the clinical findings of the nuclear YAP observed in some patient samples, implying that the detection of nuclear or cytoplasmic YAP is likely reflecting the degree of disruption in junction integrity, mediated by PV IgG.

Likewise, treatment of cells with the pathogenic monoclonal antibody AK23 also resulted in increased nuclear p53 expression, as well as cellular p21^WAF1/CIP1^ in a time and dose-dependent manner, as compared to the respective controls (Fig EV9A, B). As expected, disruption of junctions was apparent in AK23 treated cells. Again, intriguingly, the membrane distribution of p53 was also detected in the AK23 treated cells, similar to those exposed to PV sera.

Taken together, these *in vitro* findings demonstrated that treatment of keratinocytes with PV autoantibodies, as well as a pathogenic antibody targeting the adhesion site of Dsg3 (Tsunoda et al., 2003), rendered marked disruption or depletion of Dsg3 from the membrane, leading to deregulation of Hippo signalling and concomitant induction of p53 expression.

## Discussion

p53 is considered as a guardian of genome and a central player in cell response to environmental stress as it controls programmed cell death (apoptosis) and cellular senescence (Purvis et al., 2012; Vousden and Lu, 2002). p53 senses DNA damage and in response it induces a transient cell cycle arrest, allowing DNA repair or, in the case of extensive damage, promoting irreversible growth arrest (senescence) or programmed cell death (apoptosis). Therefore, p53 is critical in tumour suppression. In this report we provide the first evidence that Dsg3 acts as anti-stress protein and a suppressor of p53 in counterbalancing its response to stress signals in keratinocytes. We show that knocking down of Dsg3 in cell culture *in vitro* or ablation of Dsg3 in mice *in vivo* causes elevated expression and stabilisation of p53 and excessive apoptosis compared to controls. This effect is more pronounced or exacerbated in cells subjected to various stresses including UVB irradiation, mechanical stretching and treatment with genotoxic drugs, in which p53 is further induced in Dsg3 knockdown cells. We demonstrate that an inverse results were observed in our gain-of-function studies in which ectopic overexpression of Dsg3 resulted in suppression of p53 and cell death. Therefore, our study uncovers a novel mechanism in response of keratinocytes to stress signals or insults daily and highlights that this Dsg3-p53 pathway is crucial in the maintenance of normal tissue integrity and homeostasis and has implication in pathogenesis of the Dsg3 associated blistering disease PV (see below).

Our study has shed light on a permissive role for Dsg3 in regulation of the Hippo pathway in keratinocytes. The Hippo signaling pathway is an important regulator in control of cell differentiation and organ growth, with its deregulation contributing to cancer development (Aqeilan, 2013; Piccolo et al., 2014). This pathway comprises a cascade of kinases with one of the key downstream transcriptional coactivators being the Yes-associated protein (YAP) (Dupont et al., 2011; Panciera et al., 2017; Piccolo et al., 2014). The mechanism that controls YAP expression and activity is mediated in part by protein phosphorylation at S127 in YAP leading to YAP degradation or its cytoplasmic accumulation via binding to 14-3-3 (Panciera et al., 2017; Piccolo et al., 2014). However, the identification of upstream regulators for the Hippo pathway remains limited and there is still a lack of information on the transmembrane proteins that serve as direct regulator of this pathway. In this report we showed that the expression of both YAP gene and protein are regulated by Dsg3 and are altered by modulating the Dsg3 levels, suggesting strongly that Dsg3 acts as an upstream regulator of YAP in keratinocytes. Significantly, we identify that Dsg3 plays a key role in regulating the localization of pYAP via a mechanism of forming a complex with it and recruiting it to the plasma membrane. Although pYAP is regarded as an inactive protein, our study identifies the potential role of pYAP at the plasma membrane in keratinocytes, a process which involved Dsg3 and pYAP that could be crucial in the maintenance of the Hippo signaling pathway during keratinocyte morphogenesis and organ growth. Marked reduction of pYAP, especially at low cell densities was demonstrated in Dsg3 knockdown cells, indicating deregulation of the Hippo pathway. Although the exact underlying molecular mechanism remains elucidated this report provides an unprecedented finding for a transmembrane protein and a member of cadherin superfamily, Dsg3 that serves as a component and a surface regulator of the Hippo pathway in keratinocytes. The Dsg3 assembly with pYAP at the plasma membrane may be required for proper onset of a terminal differentiation program in keratinocytes. Indeed, we observed isochronous expression profiles of pYAP and Dsg3 during the course of keratinocyte maturation and ageing *in vitro*. Furthermore, our study also has established a key role of YAP in negative regulation of p53 (antagonistic relationship), downstream of Dsg3, in keratinocytes. YAP is a highly dynamic protein and its function in positively or negatively regulating cell growth is cell context-dependent (Piccolo et al., 2014). Here we showed that the YAP action in keratinocytes seems to promote cell proliferation as its depletion caused p53 accumulation and nuclear localization accompanied with accelerated transcription activity in cultures of low density. We showed that introduction of YAP gene in Dsg3 depleted cells was able to rescue the p53 nuclear accumulation induced by UV irradiation, and to suppress its transcription activity. Therefore, this report has uncovered the existence of a pathway containing three key components, i.e. Dsg3, YAP and p53 and suggests Dsg3 activates YAP that in turn suppresses p53 (Dsg3->YAP-Hippo-|p53). It is worth noting that this regulation pathway is specific for Dsg3 as neither E-cadherin nor desmoplakin knockdown induced similar effect to Dsg3 depletion.

The autoimmune blistering skin disease PV is characterised to be caused by IgG autoantibodies against desmosomal cadherins and that the intercellular junction homeostasis/status is altered following PV-IgG binding to Dsg3, but the precise mechanism is still controversial. Evidence of apoptosis in PV has been reported in the literature, although it has been hypothesised that the apoptosis is associated with surface receptors other than Dsg (Grando et al., 2009; Grando, 2012; Spindler et al., 2018), whereas, our study provides, for the first time, a direct link between Dsg3 and p53 in PV and suggests that antibodies targeting Dsg3 induce p53 expression and apoptosis. Thus, disruption of intercellular junctions does not solely result from steric hindrance but rather is caused by activation of Dsg3-p53 pathway via antibody binding to Dsg3. Moreover, we have demonstrated the direct involvement of deregulation of the Hippo pathway that governing tissue homeostasis, in this devastating disease. We showed that the expression and localisation of YAP are altered in PV in vivo as well as in cells treated with PV sera or the pathogenic antibody targeting Dsg3 *in vitro* due to disruption in the integrity and homeostasis of cell-cell junctions. As a consequence, accumulation of YAP and p53 was detected in cells surrounding blisters in PV or treated with PV sera. In line with these findings, elevated expression of p53 and apoptosis was also shown in the skin of Dsg3 null mice compared to Dsg3+/- control littermates. The Hippo pathway is crucial in contact inhibition of cell proliferation which is regarded as a classical paradigm of epithelial biology. Hence, our study provides first evidence of a novel mechanism by which the disruption of the Dsg3->YAP-Hippo-|p53 pathway in keratinocytes may directly be attributing to the pathophysiology and pemphigus acantholysis in PV.

In summary, the central balance between cellular proliferation and differentiation involves two fundamental pathways, p53 and Hippo signalling (Aqeilan, 2013; Kastenhuber and Lowe, 2017). This present study provides novel evidence that the desmosomal cadherin Dsg3 functions as a key upstream regulator for the Hippo signalling pathway. This, in turn, counteracts the p53 response to various stress signals in keratinocytes, establishing a crosstalk between the Hippo and p53 pathways via Dsg3, an interaction which has important implications for the pathogenesis of blister formation in PV.

## Methods

### Cell lines, animal and clinical patient samples

Various epithelial cell lines derived from skin and other tissues were used in the study, i.e. NTERT immortalised skin keratinocytes (wild-type p53: wtp53) maintained in keratinocytes serum free medium (KSFM) (17005042, Thermo Scientific); T8 cutaneous squamous cell carcinoma cell line with a frameshift mutation at amino acid 91 of *TP53* resulting in a truncated protein and making it essentially p53 null (gift from Prof. Catherine Harwood), and they were cultured in complete keratinocyte growth medium (KGM containing Dulbecco’s Modified eagle Medium (DMEM) (12-604F, Lonza):Ham’s 12 (11765054, Thermo Scientific) in the ratio of 3:1 supplemented with 10% fetal calf serum (FCS) (Biosera), epidermal growth factor (EGF) (13247-051, Invitrogen), Insulin human solution (19278, Sigma), cholera toxin (C8052, Sigma), and hydrocortisone (H4001, Sigma). MDCK (Madin Darby canine kidney) cells (wt p53) are the simple epithelial cell line, which are derived from canine kidney tubule epithelium; A431 cell line (mutant p53-R273H) is derived from vulva squamous cell carcinoma; A2780 ovarian cancer cell line (wt p53) and HCT116 colorectal carcinoma cell line (wt p53). All these cell lines were maintained in DMEM (12-604F, Lonza) supplemented with 10% FCS (Biosera, UK). Due to the low levels of endogenous Dsg3 expression these cell lines were used for the gain-of-function studies by transduction of retroviral construct pBABE.hDsg3.myc along with the empty vector control (Moftah et al., 2016; Tsang et al., 2010), namely FL Dsg3 and Vect Ct cells, respectively (Tsang et al., 2010). Cells were incubated at 37°C in a humidified atmosphere of 95% air and 5% CO_2_. The medium was changed on alternate days and cells were subjected to subculture routinely once they reached to about 70-80% confluence.

Mouse back skin samples from Dsg3 null (Dsg3-/-) and heterozygous control littermates (Dsg3-/-) were obtained as described previously (Hunefeld et al., 2018). PV sera (anonymous, 17 cases) were received from our collaborator based in First Department of Dermatovenerology, St. Anne’s Faculty Hospital, Brno, Czech Republic, and oral tissue samples of PV patients (25 PV cases and 10 normal health tissue control as well as 3 cancer patient samples) were obtained from our collaborator in Guiyang Medical University, China; all with informed patient consent and ethical approval.

### Antibodies

The following mouse (m) and rabbit (r) monoclonal/polyclonal antibodies (Abs) were used: Dsg3 mAb against the N-terminus (5H10) (sc-23912, Santa Cruz); Dsg3 rAb against the C-terminus (H145) (sc-20116, Santa Cruz); p53 mAb (DO-1) (ab1101, Abcam); p53 rAb (C-19) (sc-1311-R, Santa Cruz); MDM2 rAb (EP16627) (ab178938, Abcam); phospho MDM2 rAb (S166) (ab131355, Abcam); p21^WAF1/CIP1^ rAb (C-19) (sc-397, Santa Cruz); Bax mAb (sc-20067, Cell Signaling); Caspase3 rAb (clone C92-605, RUO) (14C10, BD Biosciences); Caspase3 rAb (9662S, Cell Signaling); active Caspase3 rAb (ab49822, Abcam); YAP rAb (D8H1X-XP, Cell Signaling); phospho YAP rAb (EP1675Y, phosphoS127) (ab76252, Abcam); Desmoplakin rAb (sc-33555, Santa Cruz); Plakoglobin mAb (PG51, Progen); Dsc2 rAb (610120, Progen); Dsg2 mAb (33-3D) was kindly received from Prof. David Garrod; E-Cadherin mAb (HECD-1) (ab1416, Abcam); Glyceraldehyde-3-phosphate dehydrogenase (GAPDH) rAb (14c10, Cell Signaling); HSC70 mAb (B6:sc-7298, Santa Cruz); β-actin mAb (8H10D10, Cell Signaling).

### siRNA and plasmid transfection and transduction

All transient siRNA transfections at 100nM final concentration were conducted either by following the protocols described previously (Mannan et al., 2011; Tsang et al., 2012a; Tsang et al., 2012b) or using DharmaFECT 1 (T-2001-02, Dharmacon) following the manufacturer’s instructions. The self-designed siRNA (AAATGCCACAGATGCAGATGA) corresponding to nucleotides 620–640 of human Dsg3 mRNA (Accession No: NM_001944) and a scrambled control sequence (AACGATGATACATGACACGAG) were synthesized by Dharmacon (USA). Other siRNA sequences for desmoplakin (Wan et al., 2007), E-cadherin (ON-TARGETplus SMARTpool L-003877-00-0005), p53 (GAAAUUUGCGUGUGGAGUA) and YAP (SMARTpool ON-TARGETplus siRNA, L-012200-00-0005) were purchased from Dharmacon. NTERT cells expressing high levels of endogenous Dsg3, were used for the loss-of-function study. In brief, 2×105 cells were seeded into 6-well plate and allowed to grow overnight. Cells were then transfected with either scrambled or specific siRNA at a final concentration of 100 nM in Opti-MEM I (31985-062, Gibco) using oligofectamine transfection reagent (12252011, Invitrogen)/ DharmaFECT 1. Transfection was performed for 4 hours before addition of FCS at a final concentration of 10%. Cells were grown overnight before being harvested with 0.25% trypsin/EDTA (T3924, Sigma) and re-plated at the densities according to the experiments.

Other cell lines including T8, HCT116, A2780, MDCK and A431 cells expressing low levels of endogenous Dsg3 were used for the gain-of-function approach. The transduction of pBABE-puro vector control and *hDsg3.myc* was generated following procedures as described previously (Moftah et al., 2016; Tsang et al., 2010). For transient transfection of plasmid DNA of pcDNA Flag YAP1 (18881, Addgene), pcDNA3.1-p53 WT and pCMV-Neo-Bam p53 R273H mutant (16439, Addgene), along with the empty vector control plasmid, cells were seeded in a 6-well plate at a density of 2×105. In following day, the transfection cocktail was prepared by mixing 2ug of plasmid with 3x of Fugene 6 reagent (E2311, Promega) in 50 ul of Opti-MEM I before adding into the culture in 1 ml of appropriate growth media. The cells were left in transfection media overnight before proceeding with further experimentation.

### Immunofluorescence staining and image analysis

All immunofluorescent staining were performed in cells seeded on coverslips. Briefly, the siRNA pre-treated or untreated cells were seeded on coverslips at various densities according to the experiments and incubated in normal keratinocyte growth medium for different time periods. For p53 staining, coverslips were fixed with ice-cold methanol for 10 minutes at room temperature and then were washed in PBS twice before staining. For YAP/pYAP staining, due to high protein solubility, 3.6% formaldehyde was used and cells were fixed for 10 minutes followed by brief treatment with 0.1% Triton X–100 in PBS for 1∼5 minutes. Immunostaining was performed following our standard protocols (Moftah et al., 2016); the nonspecific binding sites were blocked for 15-30 minutes with 10% goat serum before the primary and then the secondary antibody incubation, each lasted for 1 hour at room temperature (RT). Coverslips were washed 3 times with washing buffer (PBS containing 0.2% Tween 20) after each antibody incubation. Finally, coverslips were counterstained with DAPI for 8–10 minutes before a last wash and then mounted on slides. Images of fluorescent staining were acquired at the same exposure for each channel, with a 40x oil objective in Leica DM4000 Epi-Fluorescence microscope or a 63x oil objective in Zeiss 710 Laser Scanning Confocal Microscope. Super-resolution microscopy was performed using Zeiss 880 Laser Scanning Confocal Microscope. Image analysis was performed with ImageJ and in Excel spreadsheets. For the quantitation of nuclear p53 staining, each image was subtracted with the binary image of corresponding DAPI channel before the total immunofluorescent intensities (IMF) were determined for the nuclear subtracted image (cytoplasmic signals) and the un-subtracted image (total signals), respectively, using the same thresholding. Finally, the nuclear signals were calculated by subtracting the cytoplasmic signals from the total signals in each field in an Excel spreadsheet before statistical analysis. Data were presented as average IMF per cells in the bar charts.

The detail procedures for immunostaining of mouse back skin were described previously (Hunefeld et al., 2018). Two mice in each group, i.e. Dsg3+/- and Dsg3-/-, were included in the study. Before sectioning, the back skin tissue was stored in DMEM at −80°C and embedded into Tissue Tec Freezing medium (Jung). 5 μm thick frozen sections were placed on Silane-Prep slides (Sigma-Aldrich Chemie GmbH, München, Germany). The sections were fixed with paraformaldehyde-lysine-periodate solution and stained with antibodies directed against p-p53 (Santa Cruz, 1:50), active caspase-3 (R&D Systems, 1:50) and p21^WAF1/CIP1^ (Abcam, 1:100). For detection, secondary antibodies (anti-goat, anti-rabbit and anti-rat) tagged to Cy3 or Alexa488 (Dianova, 1:500) were used. Nuclei were counterstained with DAPI (Sigma). Images were acquired with Zeiss LSM 800 confocal laser microscope (40x and 63x objectives). For counting of positive hair follicles in the back skin of Dsg3^-/-^ and Dsg3^+/-^ mice, multiple images for each follicle were acquired using a 5x objective and then assembled in Adobe Photoshop CS6 (Adobe). Hair follicles with positive cells for p53/p21^WAF1/CIP1^ or p53/caspase3 were marked and scored. Statistical analysis was obtained by two-sided (two-tailed) Fisher’s exact test.

### Immunohistochemistry in PV specimens

Oral tissue samples from 25 PV patients and 10 normal individuals as well as 3 oral cancer patients were analysed by immunohistochemistry with the mouse anti-p53 (ZM-0408) and rabbit anti-YAP (ab52771) antibodies. Paraffin embedded tissue sections were deparaffinized, hydrated, and heated in EDTA based antigen retrieval solution (pH=8). After washing 3 times in PBS, tissue sections were subjected to incubation in 3% H2O2 in PBS for 10 minutes at room temperature followed by three washes in PBS. Then slides were incubated with the primary antibody for p53 or YAP (1:100 dilution) at 4°C overnight. The next day, sections were incubated for additional 30min at 37°C before washing 3× followed by incubation with the secondary antibody (PV-6002) for 30min at 37°C. Finally, the antibody binding was detected by incubating the tissue slides in a solution of DAB before mounting. Immunohistochemical positivity was evaluated by two independent pathologists using scoring criteria. Each section was scored by counting the positive among 100 cells per field with a high-power objective; 5 arbitrary fields were selected for scoring. According to the percentage ranges of positivity, the frequencies of expression were categorized into five grades: 0: <5% positive cells; 1: 5∼25% positive cells; 2: 25∼50% positive cells; 3: 50∼75% positive cells; 4: >75% positive cells. The staining intensity was categorized into four grades: 0: nil staining; 1: weak yellow staining; 2: staining in yellowish-brown; and 3: brown. The final score was calculated based on above two categories and thus, scores 0-1 were defined as negative and scores ≥2 were defined as positive.

### Mechanical stretch

The regimen for cyclic strain was adapted from a previous publication (Russell et al., 2004). Briefly, cells were plated and grown for 1∼2 days on collagen-coated BioFlex 6-well culture plates (Flexcell ® International corporation) with flexible silicone elastomer bottom. Each plate was placed over the loading station containing 6 planar faced posts. Cell monolayers were subjected to equiaxial cyclic mechanical stretching with strain range of amplitude in 20% and a frequency of 1 Hz, in a Flexcell FX-5000 Tension System (Flexcell International, Burlingtion, NC) for 4 hours. Cells not to be stretched were seeded in the same BioFlex plates along with the strained cells but maintained at static state without any exposure to mechanical stretch, i.e. the wells were isolated by using FlexStops (Flexcell International Corporation). Lysates were extracted either immediately after strain or transferred to static state in an incubator and harvested later for the indicated time points, before analysis by Western blotting.

### Western blotting analysis and Co-Immunoprecipitation assay (Co-IP)

Western blotting analysis and immunoprecipitation assay were conducted following procedures previously described (Brown et al., 2014; Moftah et al., 2016; Tsang et al., 2010; Tsang et al., 2012b). Briefly, cell extraction was performed to isolate proteins from cultures at approximately 90% confluence. Culture were washed with ice-cold PBS and lysed on ice with 2x sodium dodecyl sulphate (SDS) laemmli sample buffer (0.5M Tris-Cl pH6.8, 4%SDS, 20% Glycerol; 10% (v/v) 2-mercaptoethanol was added after protein assay). DC protein assay (Bio-Rad, Hertfordshire, UK) was used to determine the protein concentration for each sample. Equal amount of proteins (10 ug) were separated by SDS-PAGE at 100-150 V, followed by transferring to nitrocellulose membrane (10600002, GE healthcare) at 30 V in an electrophoresis apparatus. The non-specific binding sites on the membrane were blocked for 20-30 min at RT in blocking buffer prepared as 5% (w/v) non-fat dry milk in TTBS (Tris buffer containing 0.1% Tween 20 (P1379. Sigma)) and the membranes were incubated with the primary antibodies against specific proteins at appropriate dilutions for each antibody, for overnight at 4°C. After three washes in TTBS, the membranes were incubated with the secondary antibodies conjugated with HRP (rabbit-A6667, mouse-A0168, Sigma) with 1:1000 dilution in blocking buffer for one hour at RT. After three washes in TTBS the membranes were subjected to the chemiluminescent solution-Amersham ECL plus Western blotting detection system (28906836, GE healthcare) following the manufacturer’s instructions. The membrane was then exposed to Amersham Hyperfilm ECL and developed in an AGFA Curix 60 developer in a dark room to detect target proteins. The band densitometry of blots were performed in ImageJ software and compared between each sample after normalization with an internal loading control. For co-IP, the same protein extraction and quantification protocol was used except that the buffer used was 1× RIPA buffer (Upstate) containing a protease inhibitor cocktail (Calbiochem). For immunoprecipitation, 1mg of protein was incubated with 5 μg of primary antibody at 4°C overnight on a rotation wheel. The following day, 1.7×10^5^ Protein G magnetic Dynabeads® (10003D, Invitrogen) was added to the Lysate: antibody solution and incubated for a further 1 hour at 4° C on a rotation wheel. The bead-bound immune precipitates were washed 4× in RIPA buffer and 1× in 1×TTBS. The resulting immune precipitates were re-suspended in 11 μl of 2× sample buffer and boiled for 2 min at 95-100°C and finally, were resolved and analysed in a same manner as described for Western blotting.

*Luciferase assay.* Cells with either Dsg3 knockdown or ectopicover expression, seeded in 24-wells were transfected with p53 luciferase reporter (Plasmid #28175: 145-pGL3ctrl-3’ UTR, Addgene) using Fugene 6 transfection reagent at 0.25ug per well and after 24h incubation, lysates were harvest prior to the luciferase assay using the Luciferase Assay System (Promega) (Brown et al., 2014). Finally, the luciferase activities were normalized against protein concentration determined by Bio-Rad DC protein assay (Bio-RadLaboratories Ltd., Hertfordshire, UK).

### FACS based Cell Viability-Caspase-3 assay

The detail of the procedures for this assay was described previously (Lee et al., 2018). Harvested cells were labelled with fixable live dead stain, Zombie NIR (Near Infra-Red) (BioLegend, UK) at RT for 15 mins, then washed in PBS/BSA and fixed in Solution A (Cal-Tag, UK) for 15 min at RT. Washed cells were permeabilised in 0.25% Triton X-100 (Sigma, UK) for 15 min at RT followed by incubation with anti-active caspase-3-BV650 (Cat. No. 564096, BD Biosciences, USA) for 20 min at RT. After wash, cells were re-suspended in PBS-400 and analysed on an ACEA Bioscience Novocyte 3000 flow cytometer. Zombie NIR was excited by the 633 nm laser and collected in the 780/60 nm detector. Caspase-3-BV650 was excited by the 405 nm laser and collected at 675/30 nm. Cells were gated on FSC vs SSC removing the small debris near the origin. Cells were then gated on a dot-plot of Caspase-3-BV650 vs Zombie NIR with a quadrant placed marking off live cells in the double negative quadrant (lower left), with Caspase-3-BV650^+ve^/Zombie NIR^-ve^ (lower right) indicating apoptotic cells and lastly with Caspase-3-BV650^+ve^/Zombie NIR^+ve^ and Caspase-3-BV650^-ve-^/Zombie NIR^+ve^ upper quadrants indicating dead cells.

### RT-qPCR

The detail procedures in reverse transcription quantitative PCR (RT-qPCR) analysis was described previously (Gemenetzidis et al., 2009). Briefly, the mRNA harvested using Dynabeads mRNA Direct kit (Invitrogen) were converted to cDNA using qPCRBIO cDNA Synthesis kit (#PB30.11-10, PCRBIO Systems, UK) and the cDNA was diluted 1:5 with RNase/DNase free water and stored at −20°C until used for qPCR. Relative gene expression qPCR were performed using qPCRBIO SyGreen Blue Mix Lo Rox (#PB20.11-50, PCRBIO Systems, UK) in the 384-well LightCycler 480 qPCR system (Roche) according to our well-established protocols (Gemenetzidis et al., 2009) which are MIQE compliant (Bustin et al., 2009). Thermocycling begins with 95°C for 30s prior to 45 cycles of amplification at 95°C for 1s, 60°C for 1s, 72°C for 6s, 76°C for 1s (data acquisition). A ‘touch-down’ annealing temperature intervention (66°C starting temperature with a step-wise reduction of 0.6°C/cycle; 8 cycles) was introduced prior to the amplification step to maximise primer specificity. Melting analysis (95°C for 30s, 65°C for 30s, 65-99°C at a ramp rate of 0.11°C/s) was performed at the end of qPCR amplification to validate single product amplification in each well. Relative quantification of mRNA transcripts was calculated based on an objective method using the second derivative maximum algorithm (Roche). All target genes were normalized using a stable reference gene (POLR2A). Samples were presented as the mean ± SEM of 3 replicates. The primers used in the study are shown in Table 1.

**Table 1.**
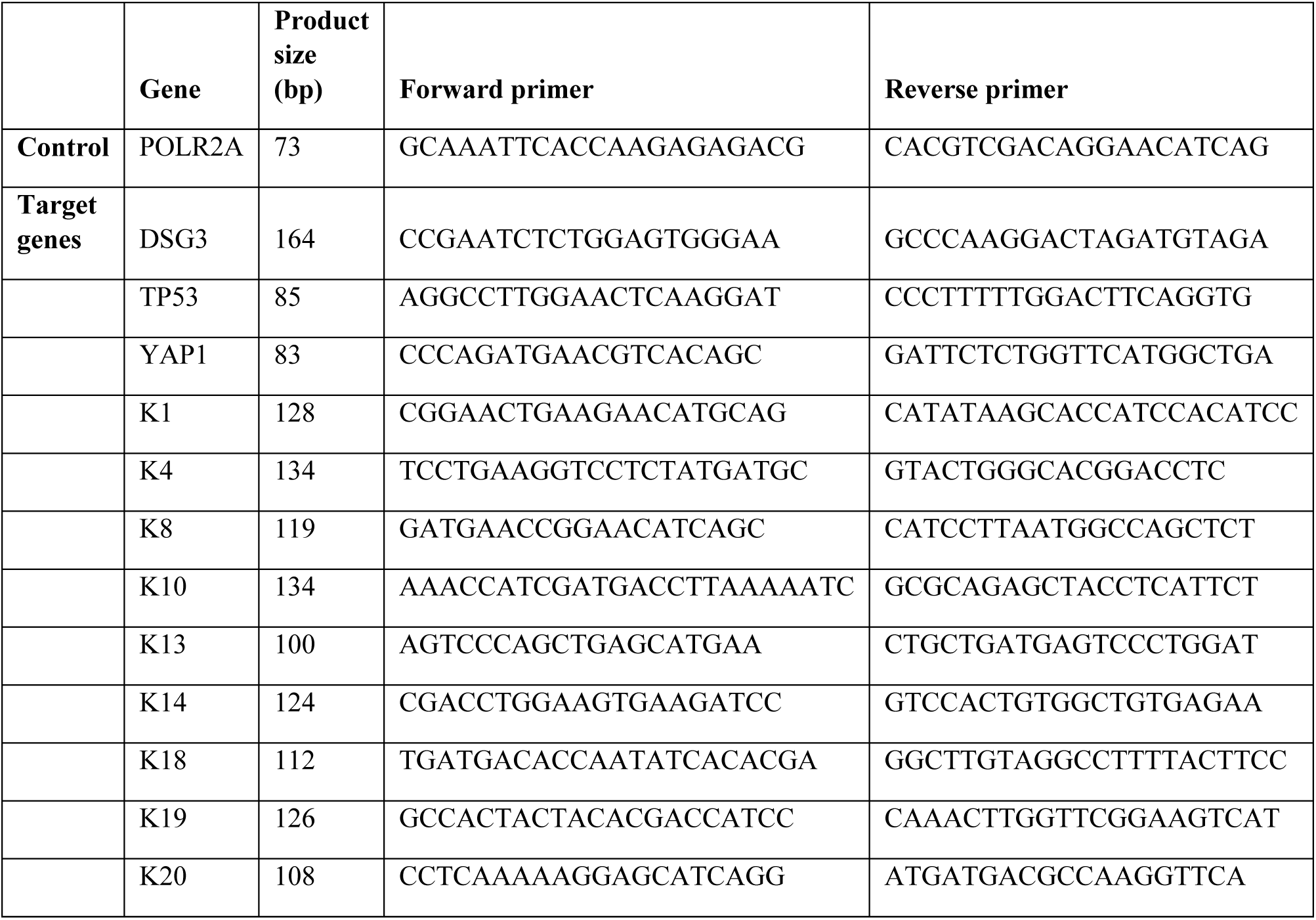
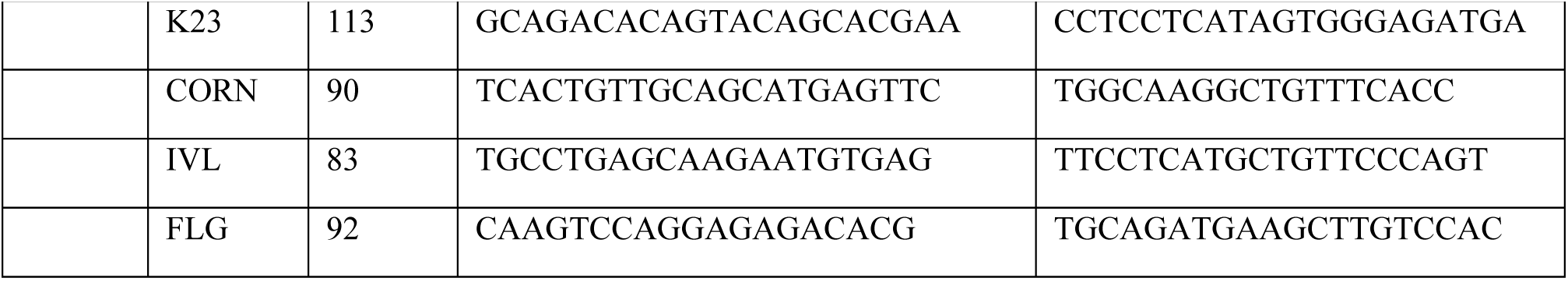
Primer sequences of keratin and terminal differentiation genes analysed in the study

### Statistical Analysis

Statistical differences between control and test groups were analysed using unpaired, 2-tailed Student *t*-test in most cases. For some experiments where sample numbers were uneven the data was additionally analysed by the Wilcoxon-Mann-Whitney Rank Test. Data are presented as mean ± s.d. unless otherwise indicated. Two-sided Fisher’s exact test was used for the comparison of the positive hair follicle scoring in mice. Chi-Square statistic was used for obtaining the p values in comparison between PV patient samples and normal healthy controls and the sample size for the clinical samples is shown in Fig. 4a,b. P values less than 0.05 was considered statistically significant, *i.e.* * p<0.05, ** p<0.01, *** p<0.001 and **** p<0.0001. Experiment was repeated at least three times. The sample size varied depending upon the individual experiments. In general, the microscopic images were acquired in 4-6 arbitrary fields per sample. For Western blotting analysis, lysates were collected from three biologically independent replicate. The represented blots were displayed in figures. Wherever possible, the comparison between control and test groups was normalised against the control and expressed as a fold change relative to the control samples that were set as 1. For the p53 protein turnover analysis, the fold change against the first time point in each condition that was set as 1, was presented.

## Expanded View

**Figure EV1.**
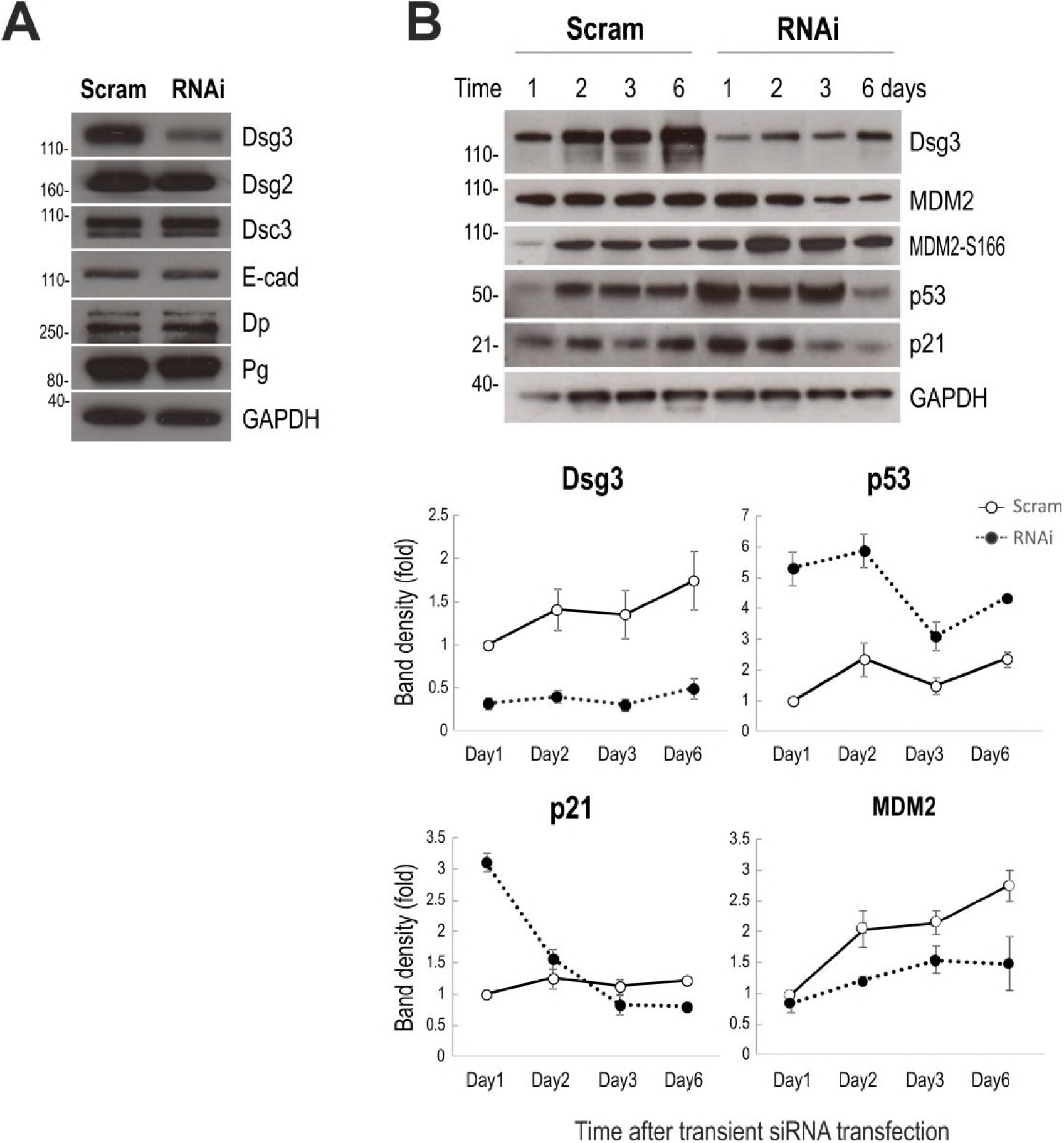
Dsg3 knockdown elicits the induction of p53 and p21^WAF1/CIP1^ as well as MDM2 in NTERT cells. (**A**) Western blots of the indicated junctional proteins in siRNA transfected cells. No evident changes were observed in other junctional proteins except for Dsg3 specific knockdown. (**B**) Time course analysis for the indicated proteins in cells with or without Dsg3 knockdown showing recovery of normal Dsg3, p53 and p21^WAF1/CIP1^ levels. The Dsg3 expression in scrambled control siRNA treated cells showed steadily increasing levels in a time-dependent manner, and reached the plateau on day 6 when the experiment was terminated, whereas p53 and p21^WAF1/CIP1^ showed small variations in contrast to that in Dsg3 knockdown cells, in which an enhanced expression in p53 and p21^WAF1/CIP1^ was detected in the first couple of days of siRNA transfection. Finally, both p53 and p21^WAF1/CIP1^ were reduced in knockdown cells by day 3∼6 when Dsg3 level was restored to some extent. Concomitantly, a decrease in MDM2 and an increase in MDM2-S166 were shown in cells with Dsg3 knockdown, compared to the respective control samples.

**Figure EV2.**
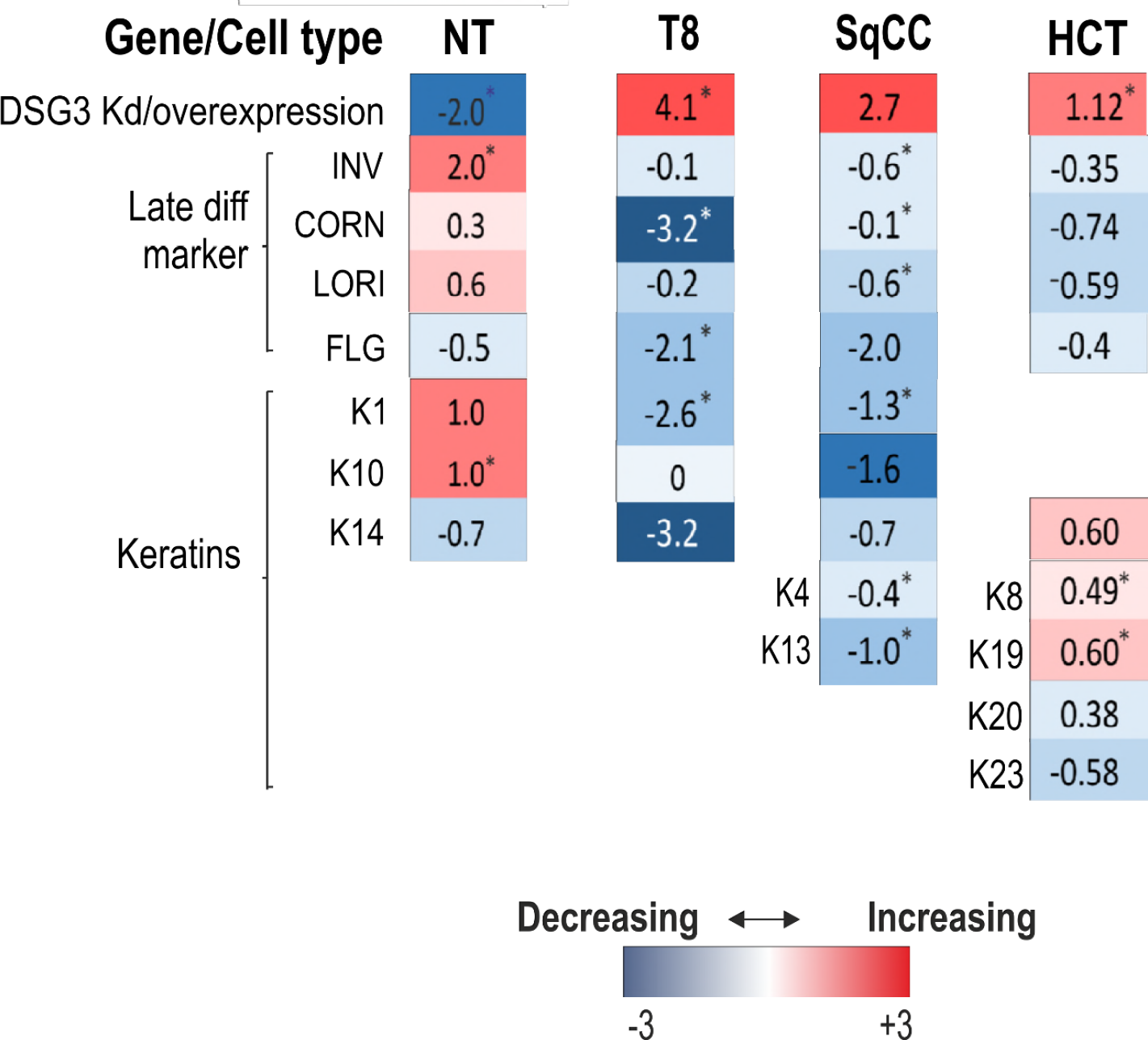
qPCR analysis in cells with either Dsg3 knockdown or ectopic overexpression indicates an inverse correlation between Dsg3 and early/late differentiation markers. Heat map of Log2 fold changes of knockdown (KD) or overexpression against to the respective controls, for the indicated genes, i.e. keratin and terminal differentiating structural genes, in various cell lines, such as NTERTs with Dsg3 knockdown, and three lines (cutaneous keratinocyte line T8, oral keratinocyte line SqCC/Y1 and colorectal HCT116 cells) with exogenous Dsg3 overexpression; n=3 biologically independent samples, asterisks indicate statistical significance via unpaired two-sided student *t*-test, with p<0.05.

**Figure EV3.**
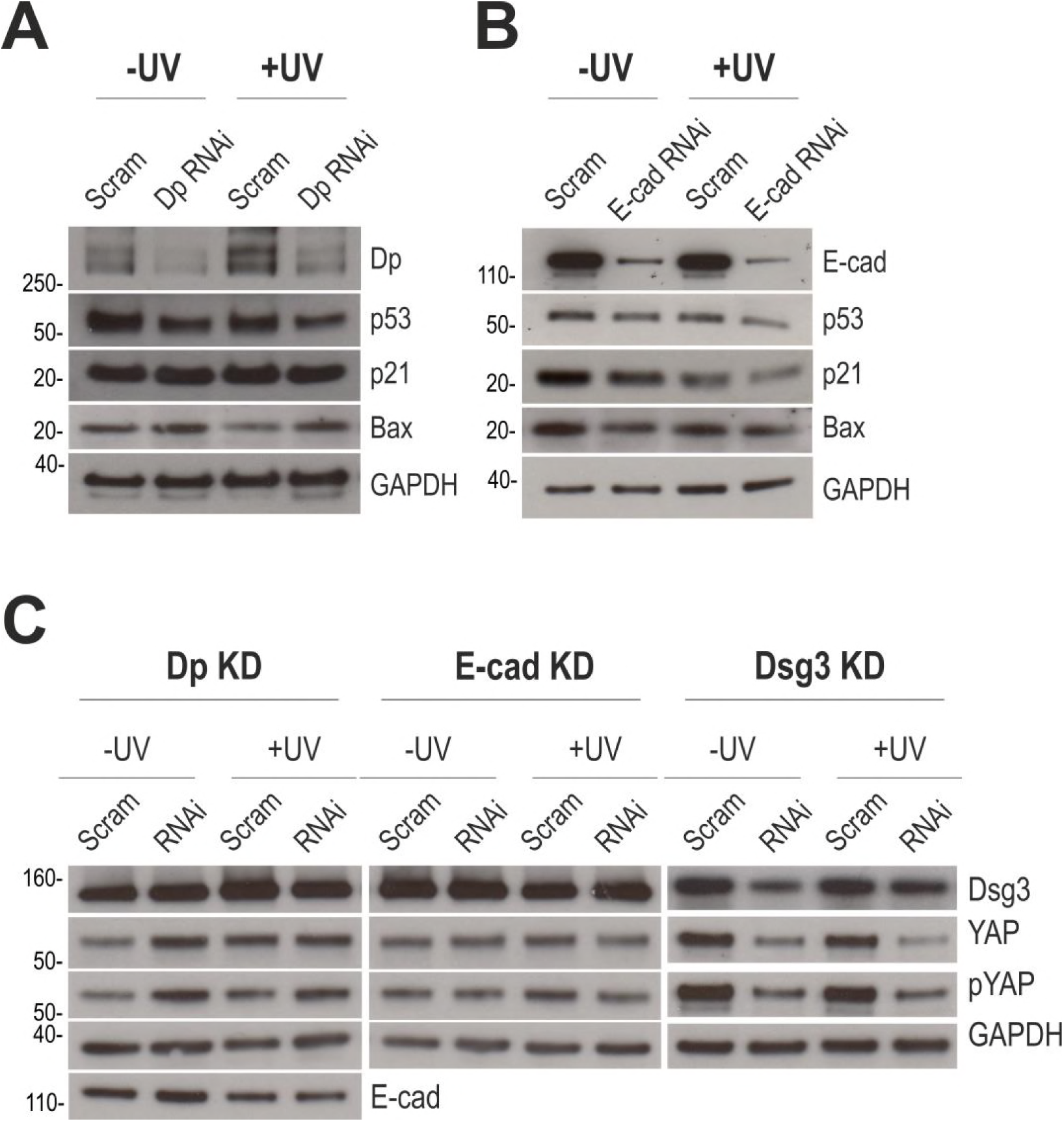
Knockdown Dp or E-cadherin in NTERT cells shows distinct protein expression profiles for p53, p21^WAF1/CIP1^, Bax, YAP and pYAP compared to cells with Dsg3 knockdown. (**A**) Western blotting analysis indicated that Dp knockdown resulted in a small reduction in p53 and Bax but little change in p21^WAF1/CIP1^ regardless UV irradiation. (**B**) E-cadherin knockdown resulted in similar changes in the expression of p53, p21^WAF1/CIP1^ and Bax that showed a small reduction compared to control cells, and UV irradiation caused a moderate further reduction compared to the respective control cells. (**C**) Different protein expression profiles were also detected for YAP and pYAP in cells with knockdown of Dp, E-cadherin and Dsg3, respectively. Regardless to UV, consistently Dsg3 knockdown resulted in a reduction of YAP and pYAP. In contrast, few changes were seen in cells with E-cadherin knockdown but a moderate increase in cells with Dp knockdown. Neither Dp nor E-cadherin knockdown had any effect on the Dsg3 expression.

**Figure EV4.**
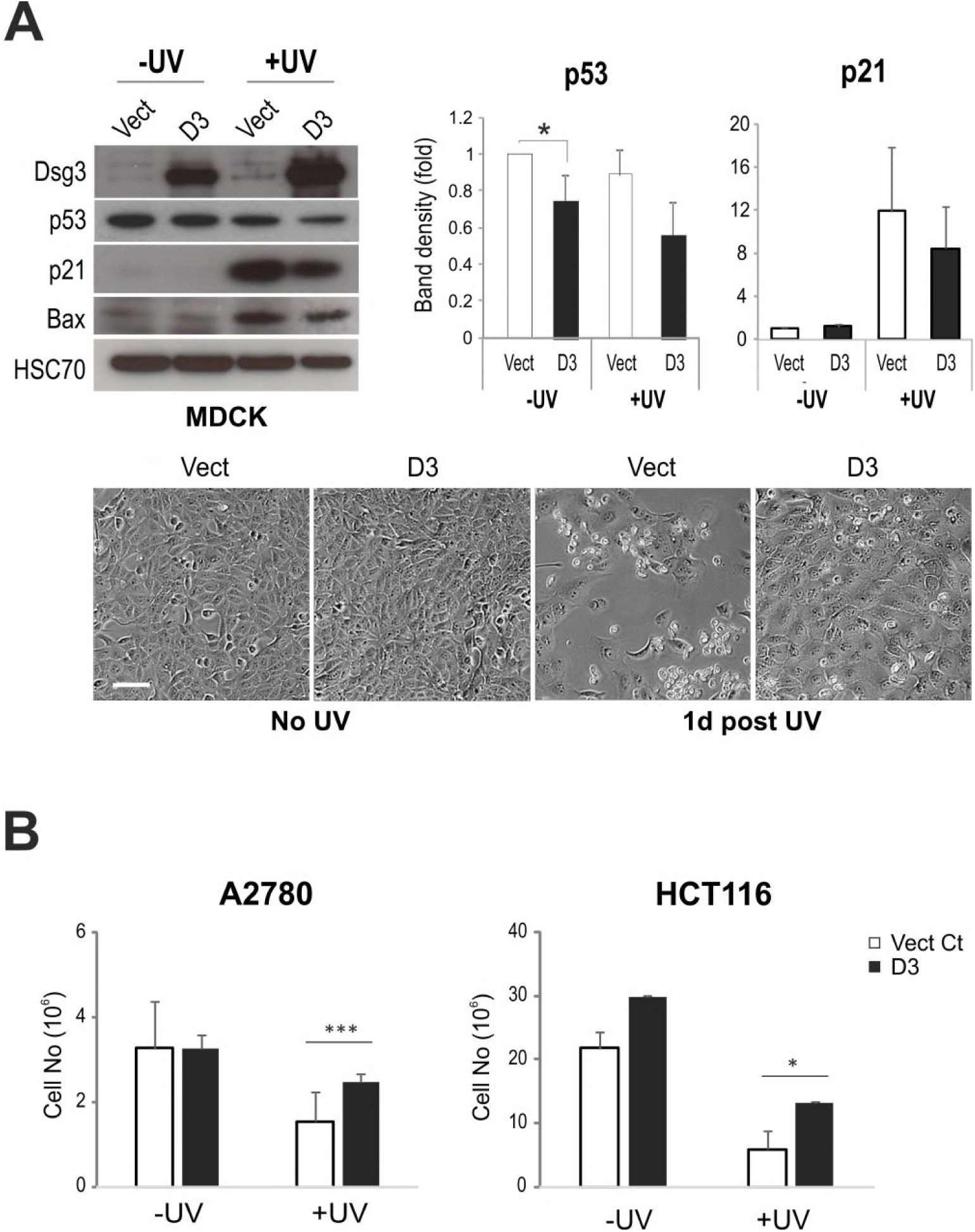
Overexpression of Dsg3 in various cell lines protects cells from the UV induced cell death. (**A**) Analysis of p53 and its targets p21^WAF1/CIP1^ and Bax in stable MDCK cell lines (harbour wtp53), Vect control and D3 with overexpression of Dsg3, without or with UV irradiation. p53/p21: n=3 biologically independent samples. The phase contrast images below showed the loss of a large sub-population of Vect cells one day after UV whereas in contrast, many D3 cells remained attached to the substrate. Scale bar, 50 μm. (**B**) A2780 (ovarry cancer line) and HCT116 (colorectal cancer line) that harbour wtp53 were seeded in 6-well plates for 1 day before treated in the absence or presnce of UVB irradiation (10mj/cm^2^). After 1 day, some UV treated cells were floating. All the attached cells in quadruplicate wells (V and D3, -/+ UV) in each condition were harvested by trypsin/EDTA and the total cell number in each well was determined by direct cell counting with a CASY machine; n=3, data are mean±s.d. Significant increased number of viable cells was detected in the Dsg3 overexpressing cell lines compared to the respective controls. All comparison were made using unpaired two-sided student *t*-test, *p<0.05, ***p<0.001.

**Figure EV5.**
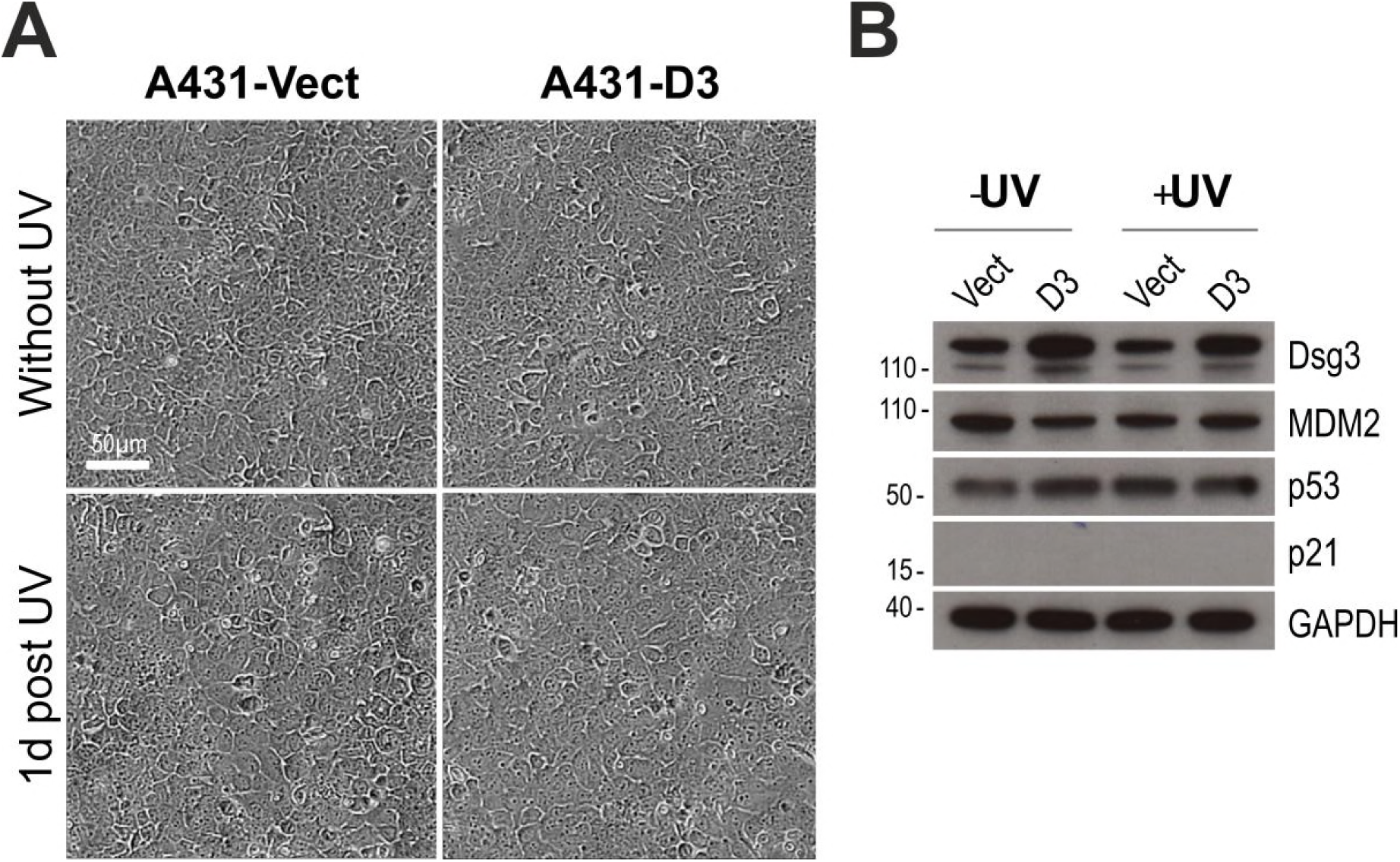
Overexpression of Dsg3 in A431 cell line showed no induction of p53 in response to UV irradiation. **a**, Phase contrast images of A431-Vect Ct and Dsg3 overexpressing (D3) lines 1 day after treatment with or without UVB (10mj/cm^2^). No any difference in cell detachment was noticeable between different samples. **b**, The representative Western blots for the indicated proteins that show no suppression in p53 in Dsg3 overexpressing cells and no detection for p21^WAF1/CIP1^ either in different samples.

**Figure EV6.**
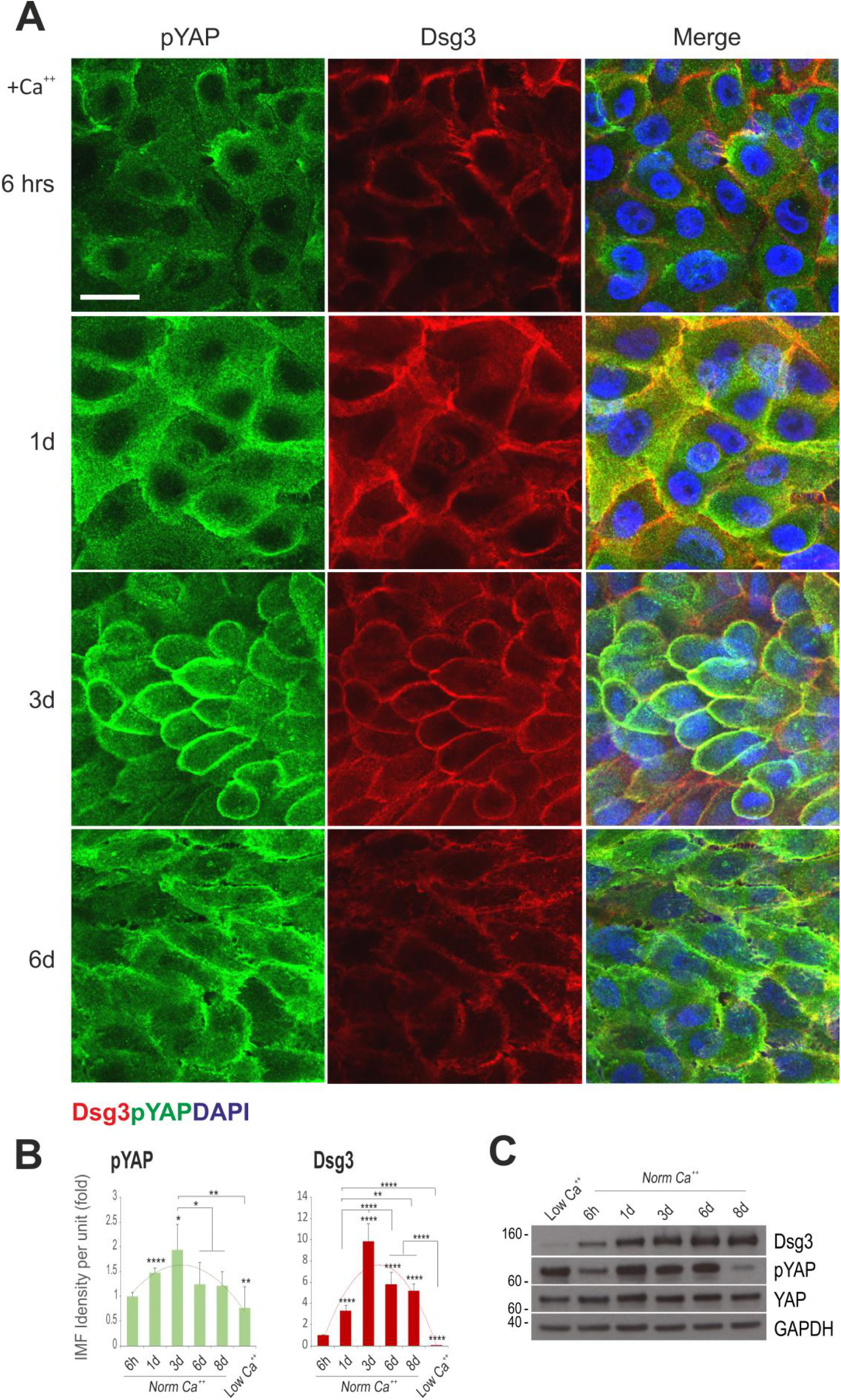
pYAP shows transient expression along with Dsg3 in confluent culture of NTERTs. (**A**) Confocal images of double immunostaining for Dsg3/pYAP in confluent NTERTs at different ages, i.e. cultures at 6 hours, 1, 3, 6 and 8 days in KGM medium containing normal calcium concentration. The low calcium medium (KSFM) treated cells were used as the control. Scale bar, 20 μm. Merged images show co-local sation of pYAP (green) and Dsg3 red) at the cell junctions (yellow). The nuclei were counterstained with DAPI. (**B**) Quantitation data of image in a; n=6 fields per condition and the data is representative of at least 4 independent experiments. *p<0.05, ***p<0.001, ****p<0.0001. Of note, the expression profiles of Dsg3 and pYAP were similar and the levels reached the highest at approximately 1-3 days before declined gradually. pYAP also exhibited distribution at the plasma membrane where it colocalised with Dsg3, especially during 1-3 days of confluent cultures. Finally, pYAP showed internalized and degraded in prolonged cultures of 6-8 days. (**C**) Western blotting analysis of YAP, pYAP and Dsg3 for the time course study shown in a&b; the representative of two independent experiments and consistent findings were observed. While the levels of total YAP showed relatively consistent during this period, the expression pYAP displayed two peaks, one in low calcium culture and another in 1 day before declined and reached to the lowest levels at day 8 when the experiment was terminated.

**Figure EV7.**
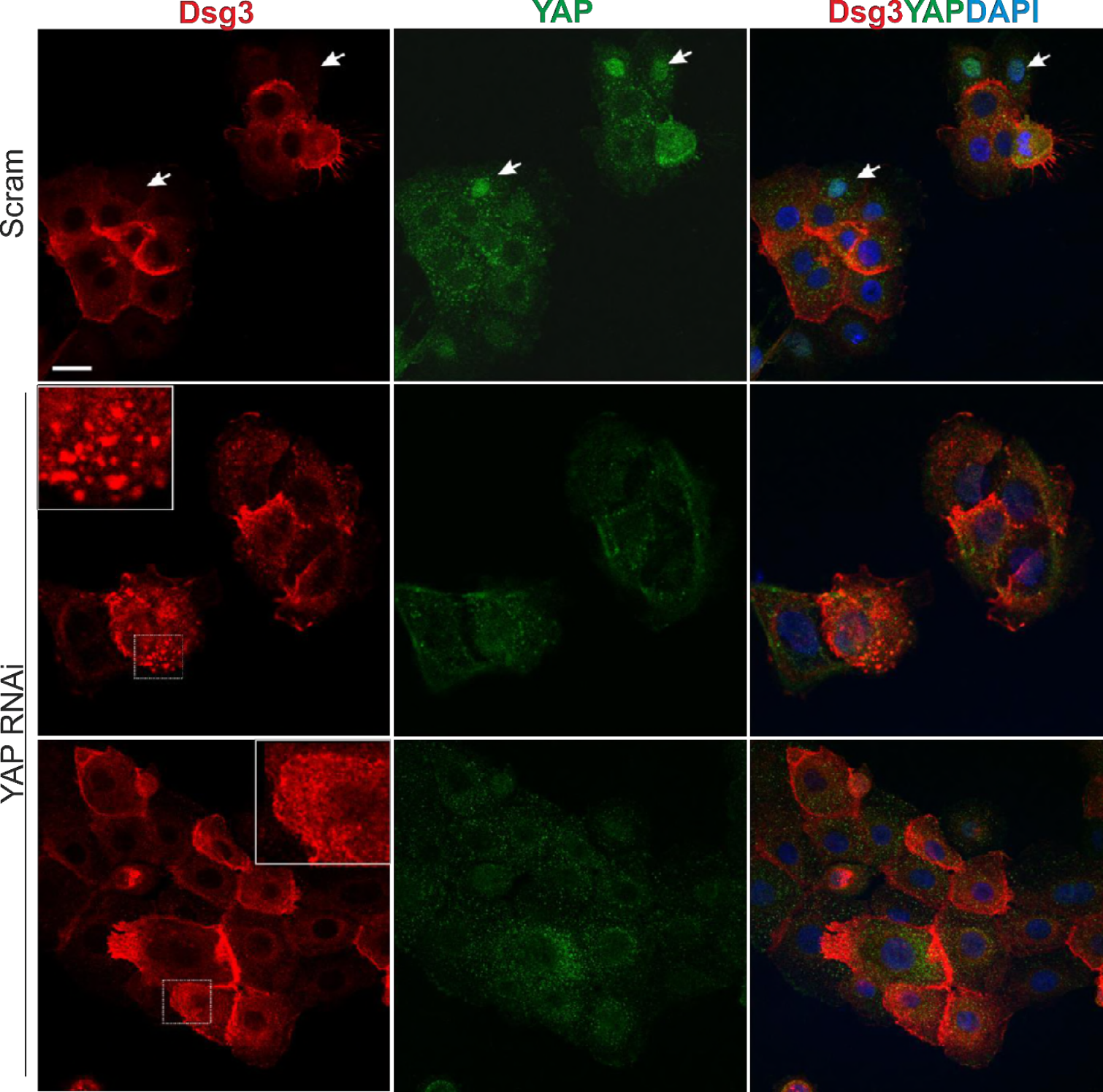
YAP knockdown caused altered and disrupted Dsg3 distribution at the junctions. Confocal images of NTERTs with or without YAP knockdown and double labelled for YAP and Dsg3. Cells were cultured in KGM with normal calcium for 6 hours before fixation. Control cells showed the Dsg3 expression predominantly located at the cell borders. In contrast, punctate expression coupled with broad diffuse distribution of Dsg3 was noticeable in cells with YAP knockdown. White arrows indicate nuclear YAP detection in control cells with relatively low levels of Dsg3 expression.

**Figure EV8.**
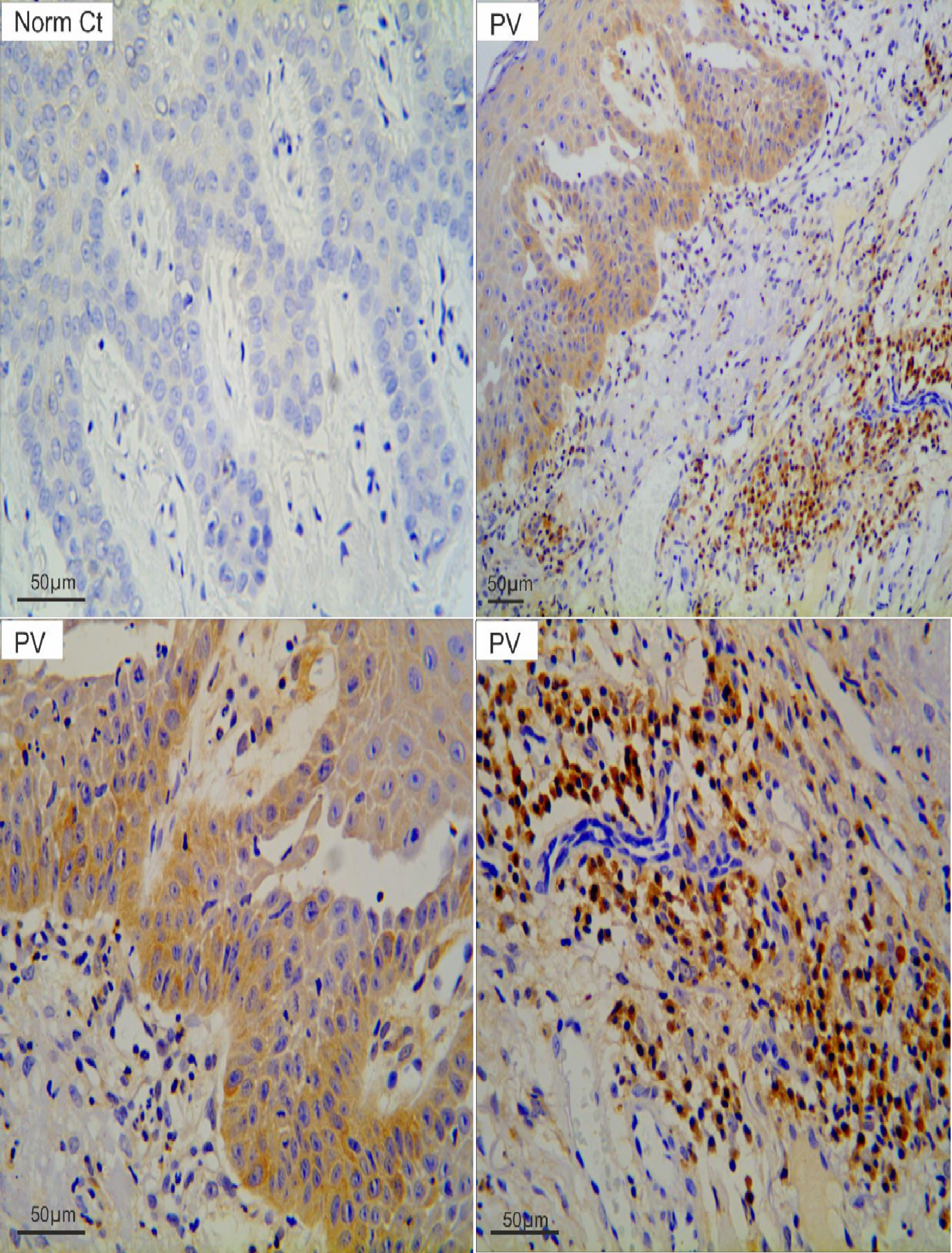
Positive staining of active caspase-3 was detected in PV patient samples. Immunohistochemistry of active caspase-3 in oral mucous tissues of normal control (top left hand panel) and PV patients (top right hand panel and bottom two panels) with the positive p53 staining also showed positive staining for cleaved caspase-3, especially in the basal and immediate suprabasal layers of stratified squamous epithelium. The positive staining was also detected in the cells located in sub-mucous connective tissue (bottom right).

**Figure EV9.**
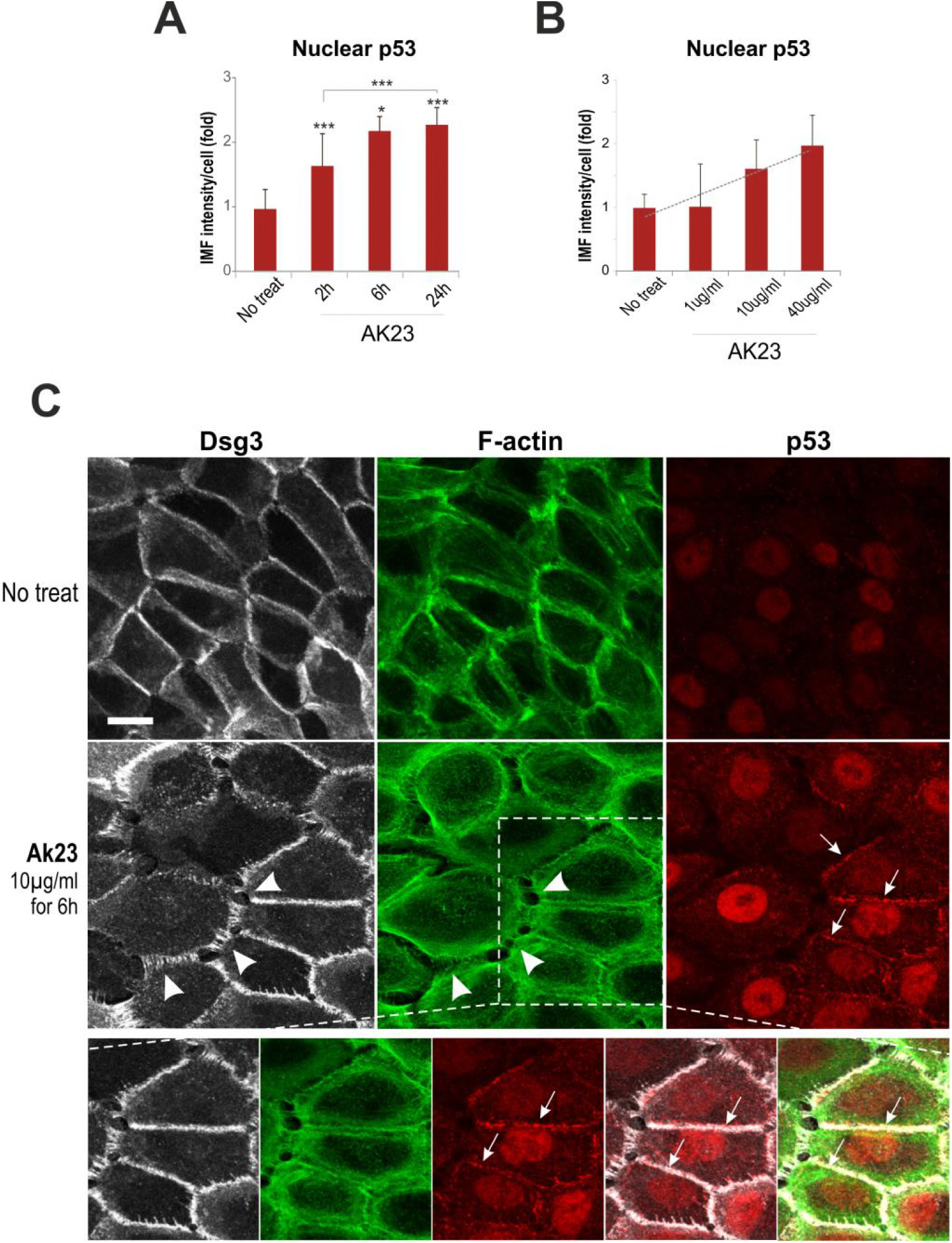
Treatment of NTERT cells with the pathogenic monoclonal antibody AK23 targeting the N-terminus of the extracellular domain in Dsg3, caused the p53 and p21^WAF1/CIP1^ induction. (**A,B**) Quantitation of immunofluorescent staining for the cellular expression of p53 in NTERT cells treated with AK23 (1 μg/ml) in a time-course, as well as dose-dependent experiments; n=11 fields per condition, pooled from 2 independent experiments. For the dose experiment, cells were treated with AK23 for 6 hours. *p<0.05, **p<0.01, ***p<0.001. (**C**) Confocal images of NTERTs with triple staining for Dsg3 (grey), F-actin (green) and p53 (red), treated in the presence and absence of AK23 (10μg/ml) for 6 hours. Disruption of F-actin along with Dsg3 (arrowheads) was readily detectable in cells exposed to AK23. Increase p53 expression was detected predominantly in the nucleus and also was observed at the plasma membrane where it showed colocalisation with Dsg3 and F-actin (arrows) in cells treated with AK23. The membrane distribution of p53 was not detectable in control cells. The protein colocalisation within the dotted line box is shown below. Scale bar, 10 μm.

## References

Amagai, M., Ahmed, A.R., Kitajima, Y., Bystryn, J.C., Milner, Y., Gniadecki, R., Hertl, M., Pincelli, C., Kurzen, H., Fridkis-Hareli, M., Aoyama, Y., Frusic-Zlotkin, M., Muller, E., David, M., Mimouni, D., Vind-Kezunovic, D., Michel, B., Mahoney, M., and Grando, S. Are desmoglein autoantibodies essential for the immunopathogenesis of pemphigus vulgaris, or just “witnesses of disease”? Exp. Dermatol. 2006; 15, 815–831

Amagai, M., Klaus-Kovtun, V., and Stanley, J.R. Autoantibodies against a novel epithelial cadherin in pemphigus vulgaris, a disease of cell adhesion. Cell. 1991; 67, 869–877

Amagai, M., Koch, P.J., Nishikawa, T., and Stanley, J.R. Pemphigus vulgaris antigen (desmoglein 3) is localized in the lower epidermis, the site of blister formation in patients. J. Invest Dermatol. 1996; 106, 351–355

Aqeilan, R.I. Hippo signaling: to die or not to die. Cell Death. Differ. 2013; 20, 1287-1288

Bar, J., Cohen-Noyman, E., Geiger, B., and Oren, M. Attenuation of the p53 response to DNA damage by high cell density. Oncogene. 2004; 23, 2128–2137

Brown, L., and Wan, H. Desmoglein 3: a help or a hindrance in cancer progression? Cancers. (Basel). 2015; 7, 266–286

Brown, L., Waseem, A., Cruz, I.N., Szary, J., Gunic, E., Mannan, T., Unadkat, M., Yang, M., Valderrama, F., O’Toole, E.A., and Wan, H. Desmoglein 3 promotes cancer cell migration and invasion by regulating activator protein 1 and protein kinase C-dependent-Ezrin activation. Oncogene. 2014; 33, 2363–2374

Bustin, S.A., Benes, V., Garson, J.A., Hellemans, J., Huggett, J., Kubista, M., Mueller, R., Nolan, T., Pfaffl, M.W., Shipley, G.L., Vandesompele, J., and Wittwer, C.T. The MIQE guidelines: minimum information for publication of quantitative real-time PCR experiments. Clin. Chem. 2009; 55, 611–622

Calkins, C.C., Setzer, S.V., Jennings, J.M., Summers, S., Tsunoda, K., Amagai, M., and Kowalczyk, A.P. Desmoglein endocytosis and desmosome disassembly are coordinated responses to pemphigus autoantibodies. J. Biol. Chem. 2006; 281, 7623–7634

Chen, Y.J., Chang, J.T., Lee, L., Wang, H.M., Liao, C.T., Chiu, C.C., Chen, P.J., and Cheng, A.J. DSG3 is overexpressed in head neck cancer and is a potential molecular target for inhibition of oncogenesis. Oncogene. 2007; 26, 467–476

Chen, Y.J., Lee, L.Y., Chao, Y.K., Chang, J.T., Lu, Y.C., Li, H.F., Chiu, C.C., Li, Y.C., Li, Y.L., Chiou, J.F., and Cheng, A.J. DSG3 facilitates cancer cell growth and invasion through the DSG3-plakoglobin-TCF/LEF-Myc/cyclin D1/MMP signaling pathway. PLoS. One. 2013; 8, e64088

Delva, E., Jennings, J.M., Calkins, C.C., Kottke, M.D., Faundez, V., and Kowalczyk, A.P. Pemphigus vulgaris IgG-induced desmoglein-3 endocytosis and desmosomal disassembly are mediated by a clathrin- and dynamin-independent mechanism. J. Biol. Chem. 2008; 283, 18303–18313

Dupont, S., Morsut, L., Aragona, M., Enzo, E., Giulitti, S., Cordenonsi, M., Zanconato, F., Le, D.J., Forcato, M., Bicciato, S., Elvassore, N., and Piccolo, S. Role of YAP/TAZ in mechanotransduction. Nature. 2011; 474, 179–183

Fang, W.K., Chen, B., Xu, X.E., Liao, L.D., Wu, Z.Y., Wu, J.Y., Shen, J., Xu, L.Y., and Li, E.M. Altered expression and localization of desmoglein 3 in esophageal squamous cell carcinoma. Acta Histochem. 2014; 116, 803–809

Ferris, R.L., Xi, L., Raja, S., Hunt, J.L., Wang, J., Gooding, W.E., Kelly, L., Ching, J., Luketich, J.D., and Godfrey, T.E. Molecular staging of cervical lymph nodes in squamous cell carcinoma of the head and neck. Cancer Res. 2005; 65, 2147–2156

Fukuoka, J., Dracheva, T., Shih, J.H., Hewitt, S.M., Fujii, T., Kishor, A., Mann, F., Shilo, K., Franks, T.J., Travis, W.D., and Jen, J. Desmoglein 3 as a prognostic factor in lung cancer. Hum. Pathol. 2007; 38, 276–283

Furth, N., Aylon, Y., and Oren, M. p53 shades of Hippo. Cell Death. Differ. 2018; 25, 81-92

Gemenetzidis, E., Bose, A., Riaz, A.M., Chaplin, T., Young, B.D., Ali, M., Sugden, D., Thurlow, J.K., Cheong, S.C., Teo, S.H., Wan, H., Waseem, A., Parkinson, E.K., Fortune, F., and Teh, M.T. FOXM1 upregulation is an early event in human squamous cell carcinoma and it is enhanced by nicotine during malignant transformation. PLoS. One. 2009; 4, e4849

Grando, S.A. Pemphigus autoimmunity: hypotheses and realities. Autoimmunity. 2012; 45, 7-35

Grando, S.A., Bystryn, J.C., Chernyavsky, A.I., Frusic-Zlotkin, M., Gniadecki, R., Lotti, R., Milner, Y., Pittelkow, M.R., and Pincelli, C. Apoptolysis: a novel mechanism of skin blistering in pemphigus vulgaris linking the apoptotic pathways to basal cell shrinkage and suprabasal acantholysis. Exp. Dermatol. 2009; 18, 764–770

Huang, C.C., Lee, T.J., Chang, P.H., Lee, Y.S., Chuang, C.C., Jhang, Y.J., Chen, Y.W., Chen, C.W., and Tsai, C.N. Desmoglein 3 is overexpressed in inverted papilloma and squamous cell carcinoma of sinonasal cavity. Laryngoscope. 2010; 120, 26–29

Hunefeld, C., Mezger, M., Muller-Hermelink, E., Schaller, M., Muller, I., Amagai, M., Handgretinger, R., and Rocken, M. Bone Marrow-Derived Stem Cells Migrate into Intraepidermal Skin Defects of a Desmoglein-3 Knockout Mouse Model but Preserve their Mesodermal Differentiation. J. Invest Dermatol. 2018; 138, 1157–1165

Kastenhuber, E.R., and Lowe, S.W. Putting p53 in Context. Cell. 2017; 170, 1062–1078

Kennedy, B.G., and Lever, J.E. Regulation of Na+,K+-ATPase activity in MDCK kidney epithelial cell cultures: role of growth state, cyclic AMP, and chemical inducers of dome formation and differentiation. - J Cell Physiol. 1984; -63

Kim, N.G., Koh, E., Chen, X., and Gumbiner, B.M. E-cadherin mediates contact inhibition of proliferation through Hippo signaling-pathway components. Proc. Natl. Acad. Sci. U. S. A. 2011; 108, 11930–11935

Kitajima, Y. 150(th) anniversary series: Desmosomes and autoimmune disease, perspective of dynamic desmosome remodeling and its impairments in pemphigus. Cell Commun. Adhes. 2014; 21, 269-280

Koch, P.J., Mahoney, M.G., Cotsarelis, G., Rothenberger, K., Lavker, R.M., and Stanley, J.R. Desmoglein 3 anchors telogen hair in the follicle. J. Cell Sci. 1998; 111 (Pt 17), 2529–2537

Koch, P.J., Mahoney, M.G., Ishikawa, H., Pulkkinen, L., Uitto, J., Shultz, L., Murphy, G.F., Whitaker-Menezes, D., and Stanley, J.R. Targeted disruption of the pemphigus vulgaris antigen (desmoglein 3) gene in mice causes loss of keratinocyte cell adhesion with a phenotype similar to pemphigus vulgaris. J. Cell Biol. 1997; 137, 1091-1102

Kubbutat, M.H., Jones, S.N., and Vousden, K.H. Regulation of p53 stability by Mdm2. Nature. 1997; 387, 299–303

Lanza, A., Cirillo, N., Femiano, F., and Gombos, F. How does acantholysis occur in pemphigus vulgaris: a critical review. J. Cutan. Pathol. 2006; 33, 401–412

Lee, H.L., Pike, R., Chong, M.H.A., Vossenkamper, A., and Warnes, G. Simultaneous flow cytometric immunophenotyping of necroptosis, apoptosis and RIP1-dependent apoptosis. Methods. 2018; 134-135, 56–66

Leighton, J., Estes, L.W., Mansukhani, S., and Brada, Z. A cell line derived from normal dog kidney (MDCK) exhibiting qualities of papillary adenocarcinoma and of renal tubular epithelium. - Cancer. 1970; -8

Lever, J.E. Inducers of mammalian cell differentiation stimulate dome formation in a differentiated kidney epithelial cell line (MDCK). Proc. Natl. Acad. Sci. U. S. A. 1979; 76, 1323-1327

Liu, Z.Q., Tian, Y.Q., Ma, F.R., Zhu, L., and Hu, Y.F. [Expression of desmoglein 3 in nasopharyngeal carcinoma: research of 22 cases]. Zhonghua Yi. Xue. Za Zhi. 2007; 87, 2541-2543

Mannan, T., Jing, S., Foroushania, S.H., Fortune, F., and Wan, H. RNAi-mediated inhibition of the desmosomal cadherin (desmoglein 3) impairs epithelial cell proliferation. Cell Prolif. 2011; 44, 301–310

Meek, D.W., and Knippschild, U. Posttranslational modification of MDM2. Mol. Cancer Res. 2003; 1, 1017–1026

Moftah, H., Dias, K., Apu, E.H., Liu, L., Uttagomol, J., Bergmeier, L., Kermorgant, S., and Wan, H. Desmoglein 3 regulates membrane trafficking of cadherins, an implication in cell-cell adhesion. Cell Adh Migr. 2016; 1-22

Oberleithner, H., Vogel, U., and Kersting, U. Madin-Darby canine kidney cells. I. Aldosterone-induced domes and their evaluation as a model system. - Pflugers Arch. 1990; - 32

Panciera, T., Azzolin, L., Cordenonsi, M., and Piccolo, S. Mechanobiology of YAP and TAZ in physiology and disease. Nat. Rev. Mol. Cell Biol. 2017; 18, 758–770

Piccolo, S., Dupont, S., and Cordenonsi, M. The biology of YAP/TAZ: hippo signaling and beyond. Physiol Rev. 2014; 94, 1287–1312

Piette, J., Neel, H., and Marechal, V. Mdm2: keeping p53 under control. Oncogene. 1997; 15, 1001–1010

Purvis, J.E., Karhohs, K.W., Mock, C., Batchelor, E., Loewer, A., and Lahav, G. p53 dynamics control cell fate. Science. 2012; 336, 1440–1444

Reuven, N., Adler, J., Meltser, V., and Shaul, Y. The Hippo pathway kinase Lats2 prevents DNA damage-induced apoptosis through inhibition of the tyrosine kinase c-Abl. Cell Death. Differ. 2013; 20, 1330–1340

Rotzer, V., Hartlieb, E., Vielmuth, F., Gliem, M., Spindler, V., and Waschke, J. E-cadherin and Src associate with extradesmosomal Dsg3 and modulate desmosome assembly and adhesion. Cell Mol. Life Sci. 2015; 72, 4885–4897

Rotzer, V., Hartlieb, E., Winkler, J., Walter, E., Schlipp, A., Sardy, M., Spindler, V., and Waschke, J. Desmoglein 3-Dependent Signaling Regulates Keratinocyte Migration and Wound Healing. J. Invest Dermatol. 2016; 136, 301–310

Russell, D., Andrews, P.D., James, J., and Lane, E.B. Mechanical stress induces profound remodelling of keratin filaments and cell junctions in epidermolysis bullosa simplex keratinocytes. J. Cell Sci. 2004; 117, 5233–5243

Savci-Heijink, C.D., Kosari, F., Aubry, M.C., Caron, B.L., Sun, Z., Yang, P., and Vasmatzis, G. The role of desmoglein-3 in the diagnosis of squamous cell carcinoma of the lung. Am. J. Pathol. 2009; 174, 1629–1637

Spindler, V., Eming, R., Schmidt, E., Amagai, M., Grando, S., Jonkman, M.F., Kowalczyk, A.P., Muller, E.J., Payne, A.S., Pincelli, C., Sinha, A.A., Sprecher, E., Zillikens, D., Hertl, M., and Waschke, J. Mechanisms Causing Loss of Keratinocyte Cohesion in Pemphigus. J. Invest Dermatol. 2018; 138, 32–37

Teh, M.T., Parkinson, E.K., Thurlow, J.K., Liu, F., Fortune, F., and Wan, H. A molecular study of desmosomes identifies a desmoglein isoform switch in head and neck squamous cell carcinoma. J Oral Pathol. Med. 2011; 40, 67–76

Tsang, S.M., Brown, L., Gadmor, H., Gammon, L., Fortune, F., Wheeler, A., and Wan, H. Desmoglein 3 acting as an upstream regulator of Rho GTPases, Rac-1/Cdc42 in the regulation of actin organisation and dynamics. Exp. Cell Res. 2012a; 318, 2269–2283

Tsang, S.M., Brown, L., Lin, K., Liu, L., Piper, K., O’Toole, E.A., Grose, R., Hart, I.R., Garrod, D.R., Fortune, F., and Wan, H. Non-junctional human desmoglein 3 acts as an upstream regulator of Src in E-cadherin adhesion, a pathway possibly involved in the pathogenesis of pemphigus vulgaris. J Pathol. 2012b; 227, 81–93

Tsang, S.M., Liu, L., Teh, M.T., Wheeler, A., Grose, R., Hart, I.R., Garrod, D.R., Fortune, F., and Wan, H. Desmoglein 3, via an interaction with E-cadherin, is associated with activation of Src. PLoS. One. 2010; 5, e14211

Tsunoda, K., Ota, T., Aoki, M., Yamada, T., Nagai, T., Nakagawa, T., Koyasu, S., Nishikawa, T., and Amagai, M. Induction of pemphigus phenotype by a mouse monoclonal antibody against the amino-terminal adhesive interface of desmoglein 3. J. Immunol. 2003; 170, 2170–2178

Vousden, K.H., and Lu, X. Live or let die: the cell’s response to p53. Nat. Rev. Cancer. 2002; 2, 594–604

Wan, H., South, A.P., and Hart, I.R. Increased keratinocyte proliferation initiated through downregulation of desmoplakin by RNA interference. Exp. Cell Res. 2007; 313, 2336–2344

